# The senescent mesothelial matrix accentuates colonization by ovarian cancer cells

**DOI:** 10.1101/2023.06.02.543239

**Authors:** Bharat Vivan Thapa, Tilmann Glimm, Deepak K Saini, Ramray Bhat

## Abstract

Ovarian cancer is amongst the most morbid of gynecological malignancies due to its diagnosis at an advanced stage, a transcoelomic mode of metastasis, and rapid transition to chemotherapeutic resistance. Like all other malignancies, the progression of ovarian cancer may be interpreted as an emergent outcome of the conflict between metastasizing cancer cells and the natural defense mounted by microenvironmental barriers to such migration. Here, we asked whether senescence in coelom-lining mesothelia, brought about by drug exposure, affects their interaction with disseminated ovarian cancer cells. We observed that cancer cells adhered faster on, senescent human and murine mesothelial monolayers than non-senescent controls. Time-lapse epifluorescent microscopy showed that mesothelial cells were cleared by a host of cancer cells that surrounded the former, even under sub-confluent conditions. A multiscale computational model predicted that such colocalized mesothelial clearance under sub-confluence requires greater adhesion between cancer cells and senescent mesothelia. Consistent with the prediction, we observed that senescent mesothelia expressed extracellular matrix with higher levels of fibronectin, laminins and hyaluronan than non-senescent controls. On senescent matrix, cancer cells adhered more efficiently, spread better, and moved faster and persistently, aiding the spread of cancer. Inhibition assays using RGD cyclopeptides suggested the adhesion was predominantly contributed by fibronectin and laminin. These findings led us to propose that the senescence-associated matrisomal phenotype of peritoneal barriers enhances the colonization of invading ovarian cancer cells and their clearance contributing to the metastatic burden associated with the disease.

## INTRODUCTION

The high morbidity associated with epithelial ovarian cancer (EOC) is due to its ability to rapidly metastasize and colonize the visceral peritoneum. Transformed cells at the primary site are shed into the peritoneal cavity via the movement of peritoneal fluid. Ovarian cancer cells adhere to the lining of the peritonea ^1, 2^, retract mesothelial cells ^3^, and actively degrade the underlying and exposed extracellular matrix by expressing proteolytic enzymes such as matrix metalloproteinases (MMPs) ^2^. Management of EOC involves surgical cytoreduction with chemotherapy provided before or/and after surgery ^4^. Cytotoxic therapeutics commonly employed in EOC are carboplatin and doxorubicin, which bind to and alkylate DNA, and paclitaxel, which binds to tubulin and prevents karyokinesis. Treatment with these agents inhibits cell division and ultimately leads to cell death. However, exposure to such drugs may also induce a cytostatic (as opposed to cytotoxic) state in both cancer and primary cells, also known as senescence, wherein the cells are metabolically active but growth-arrested. Therapy-induced senescence (TIS) has been proposed to suppress tumor cell proliferation ^5^. On the other hand, it has also been shown to potentiate tumor heterogeneity, resistance ^6^, and relapse ^7^. For instance, the TIS of ovarian cancer cells leads to their transient dormancy, which can be reversed to re-initiate proliferation ^8, 9^. TIS-associated cytokine secretion (as part of the broader phenomenon of senescence-associated secretory phenotype or SASP) may accentuate infiltration by tumor-associated macrophages (TAMs) in primary tumors leading to greater invasion and metastasis ^10–12^.

Exposure to therapeutic drugs can affect not just the transformed cell niche within a tumor but also the untransformed cellular microenvironment in ways that engender disease progression. Stromal constituents such as endothelial cells and bone marrow-derived cells signal in response to chemotherapeutic exposure to accentuate metastasis ^13–16^. In this context, it is pertinent to study whether senescence in stromal constituents due to drug exposure contributes to cancer cell invasion. Indeed, the accumulation of senescent/aged mesothelia, the cells lining the peritoneal cavity, has been shown to aid ovarian cancer metastasis ^17^.

In this study, we sought to ask whether chemotherapeutic drugs can drive mesothelial cells to senescence with an unintended potentiation of cancer progression. Using a human-murine hybrid ex vivo system, human mesothelial-cancer coculture system and a Cellular Potts Model (CPM)-based multiscale framework, we found that the senescent mesothelia secrete a unique matrisome that acts as a habitable niche for metastasizing cancer cells. A comprehensive understanding of how such a ‘senescence-associated matrisomal phenotype’ leads to higher invasion by ovarian cancer will help design novel therapeutics for the management of metastasis.

## RESULTS

### Senescent mesothelial monolayers potentiate adhesion of ovarian cancer cells

Parietal peritoneal tissue was dissected from 4-6 week Balb/c female mice and cultivated ex vivo as described earlier^18^. Senescence was induced in the explants using 100 nM doxorubicin. Mesothelial cells were harvested from control and senescent murine peritonea and senescence was confirmed by morphological examination (Figure S1A). Control and senescent explants were cultivated with a suspension of GFP-labelled SKOV-3 cells and the adhesion of the latter assessed (Figure 1A). SKOV-3 cells were observed to attach faster to the senescent murine peritoneum than untreated controls (Figure 1Bi and 1Bii; statistical significance shown in Figure 1Biii). To confirm this observation, an immportalized human mesothelial cell line MeT-5A was made senescent using 100 nM doxorubicin. The senescent mesothelial cells showed enlarged, flattened, and fragmented nuclear morphology (Figure S2A). The senescent MeT-5A monolayers stained positive for SA-β-Gal staining, the most common marker for senescence (Figure S2B). We also observed higher expression of the molecular marker p21 in senescent MeT-5A monolayers (Figure S2C). An adhesion assay similar to that in Figure 1B was performed on RFP-labelled senescent MeT-5A monolayers (Figure 1C). We observed significantly higher adhesion of GFP-labelled SKOV-3 cells on senescent MeT-5A monolayers (Figure 1Di, 1Dii, and 1Diii). This effect was further validated by performing the adhesion assay on senescent MeT-5A monolayers using GFP-labelled OVCAR-3 cells (Figure S3A). We observed significantly higher adhesion of OVCAR-3 cells on senescent MeT-5A monolayers than on control MeT-5A (Figure S3B).

**Figure 1:**
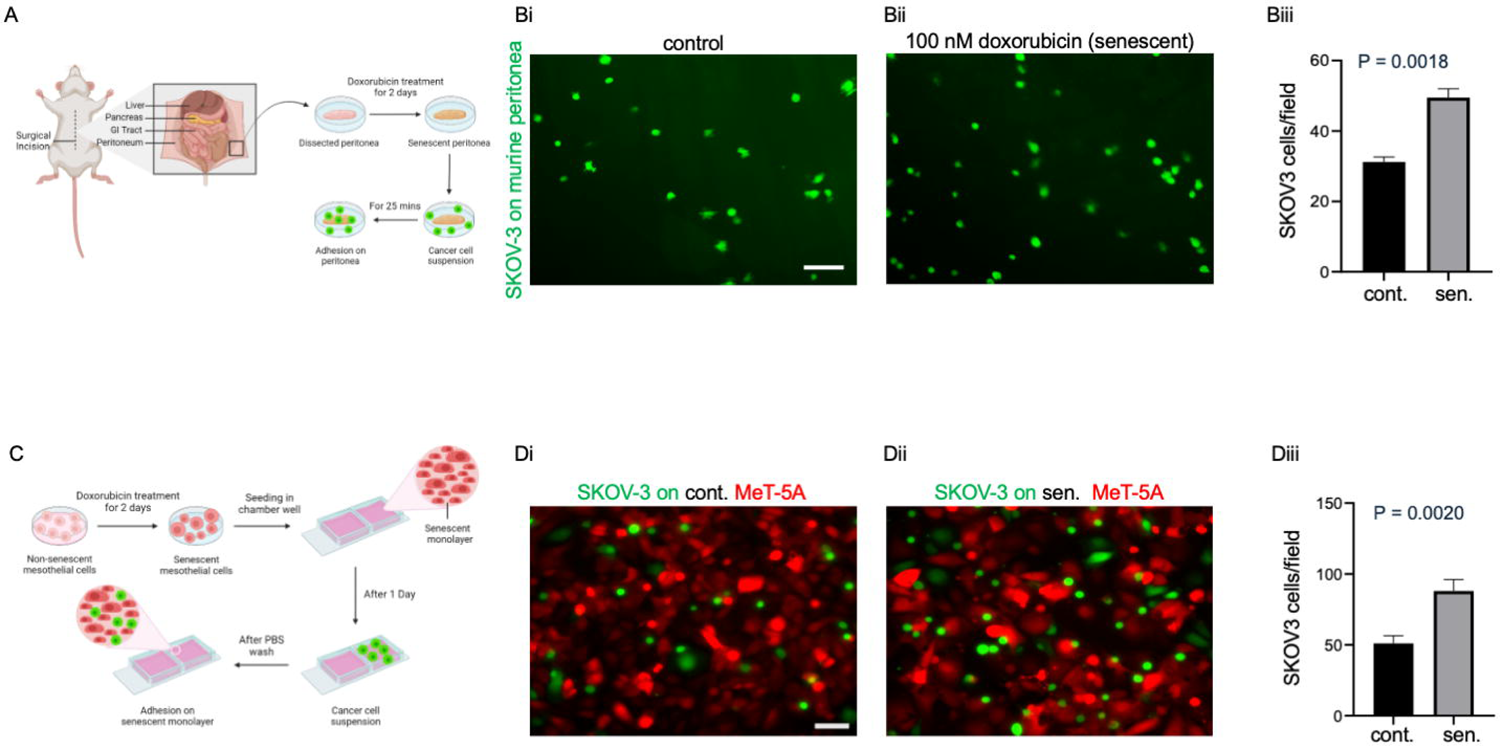
Effect of a senescent mesothelial microenvironment on the adhesion of ovarian cancer cells. (A) Schematic depiction of the adhesion assay performed on murine peritonea that were rendered senescent using 100 nM doxorubicin, cultivated ex-vivo in the presence of ovarian cancer cell suspension. (B) Representative fluorescence micrographs showing the adhesion of GFP-labelled SKOV-3 cells on control murine peritonea (i), senescent murine peritonea (ii), and bar graph showing the number of adhered SKOV-3 cells per field for control and senescent murine peritonea (iii). (C) Schematic depiction of the adhesion assay performed on RFP-labelled control and doxorubicin-induced senescent MeT-5A monolayers cultivated in the presence of GFP-labelled ovarian cancer cells. (D) Representative fluorescence micrographs showing the adhesion of GFP-labelled SKOV-3 cells on control MeT-5A monolayer (i), senescent MeT-5A monolayer (ii), and a bar graph showing the number of SKOV-3 cells per field for control and senescent MeT-5A monolayers (iii). The fields are at 10x magnification with a scale bar of 100 μm. The experiments were performed in triplicates. The data are presented as mean +/- SEM and significance is obtained using an unpaired parametric t-test with Welch’s correction.

### Ovarian cancer cells proliferate faster within a senescent mesothelial microenvironment

We asked if ovarian cancer cells not just adhered but managed to grow better in a mesothelial microenvironment with therapy-induced senescence. To investigate this, we performed time-lapse epifluorescent microscopy on monolayers of RFP-expressing control and senescence-induced MeT-5A mesothelia cultured with suspended GFP-expressing SKOV-3 cells (Figures 2A and 2B; can be seen in videos S1 and S2, respectively). We observed that the adhered SKOV-3 cells could not only proliferate better within the senescent MeT-5A monolayers (Figure 2Ci) but also clear senescent monolayers more efficiently (Figure 2Di). After 72 hours of cancer cell addition, we found a significantly higher area occupied by SKOV-3 cells within the senescent MeT-5A monolayers compared to the area occupied by SKOV-3 cells within control MeT-5A monolayers (Figure 2Cii). We also observed a statistically significant reduction in the area of senescent MeT-5A monolayers compared to control MeT-5A monolayers for the same fields (Figure 2Dii). We did not observe any notable difference in acellular spaces between control and senescent mesothelial monolayers prior to the adhesion of cancer cells, which suggested that clearance rather than available adherable space could play a role in a greater cancer cell occupancy within senescent cell populations.

**Figure 2:**
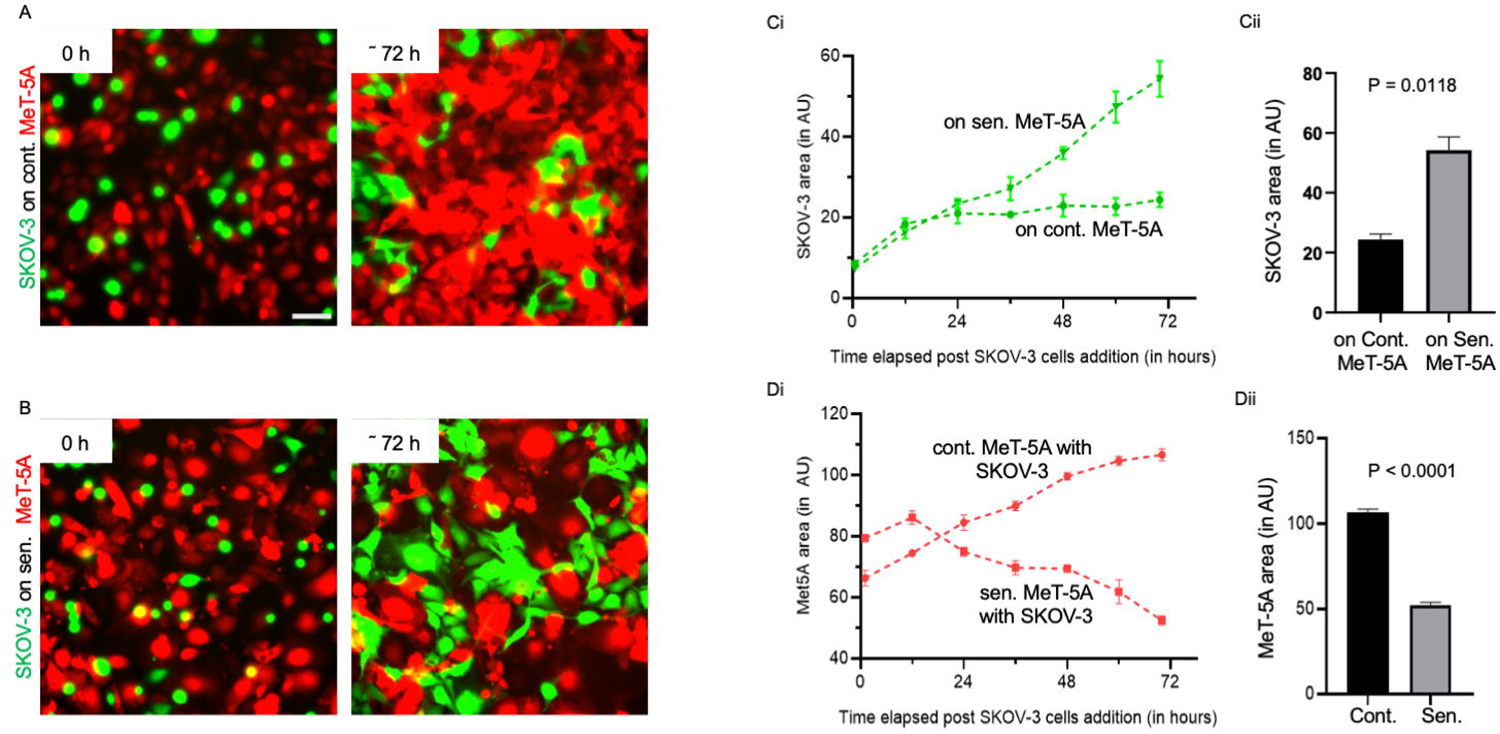
Ovarian cancer cells invade by clearing mesothelial cells. (A) Representative epifluorescence micrgraphs showing the area of control MeT-5A monolayer and SKOV-3 cells at 0 hour and 72 hour time points (from video S1). (B) Representative epifluorescent micrographs showing the area of senescent MeT-5A monolayer and SKOV-3 cells at 0 hour and 72 hour time points (from video S2). (C) (i) Quantification of the area occupied by SKOV-3 cells within control and senescent MeT-5A monolayers for 72 hours, plotted with 12-hour intervals. (ii) The bar graph compares the area of SKOV-3 cells in the two monolayers at a 72-hour time point. (D) (i) Quantification of the area of control and senescent MeT-5A monolayers for 72 hours, plotted with 12 hour intervals. (ii) The bar graph compares the area of control and senescent monolayers at a 72 hour time point. The fields are at 10x magnification with a scale bar of 100 μm. The experiments were performed in triplicates. The data are presented as mean +/- SEM and significance is obtained using an unpaired parametric t-test with Welch’s correction.

To confirm that the clearance of senescent MeT-5A monolayers is indeed caused by SKOV-3 cells, a senescent monolayer without SKOV-3 cells was also imaged for 72 hours (Figure S4A and Figure S4B; can be seen in videos S2 and S3). We observed a significant reduction in the area occupied by senescent MeT-5A monolayers in the presence of SKOV-3 cells compared to SKOV-3-less controls (Figure S4C). To confirm the ability of ovarian cancer cells to clear and proliferate amid senescent mesothelia, GFP-labelled OVCAR-3 cells were cultivated with senescent and control MeT-5A monolayers for 60 hours (Figure S5A and S5B). We observed higher proliferation of OVCAR-3 cells within senescent MeT-5A monolayers (Figure S5Ci and S5Cii) and a significant reduction in the area occupied by senescent MeT-5A monolayers compared with control MeT-5A monolayers (Figure S5Di and S5Dii). We also observed a substantial decrease in the area occupied by senescent MeT-5A monolayers cultured with OVCAR-3 cancer cells compared with senescent MeT-5A monolayers where OVCAR-3 cells were not added (Figure S6B and S6C).

### Multiple cancer cells surround and clear senescent mesothelia independent of confluence

Extrusion of cells within populations is well known across developmental and oncological contexts and is often driven through cell-density dependent mechanisms ^19, 20^. We asked if the clearance of mesothelia was achieved by cancer cells only when both were part of a jammed confluent population. To answer this question, we examined our epifluorescence time-lapses and observed senescent cell extrusions by SKOV-3 cells even in sub-confluent conditions (Figure 3A). We found that the total number of extrusions in senescent MeT-5A monolayers with SKOV-3 cells was significantly higher than the total number of extrusions in control MeT-5A monolayers with SKOV-3 cells and senescent MeT-5A monolayers without SKOV-3 cells (Figure 3B). Moreover, we observed that a significant number of extruded senescent MeT- 5A were surrounded by and impacted by more than one SKOV-3 cell before their clearance (Figure 3C). To our surprise, we also observed that senescent cells distant from SKOV-3 cells showed a lower frequency for extrusion (similar to non-senescent cells in controls distant from SKOV-3 cells. This suggested that the extrusion of senescent mesothelial cells was strongly correlated with their proximity with ovarian cancer cells.

**Figure 3:**
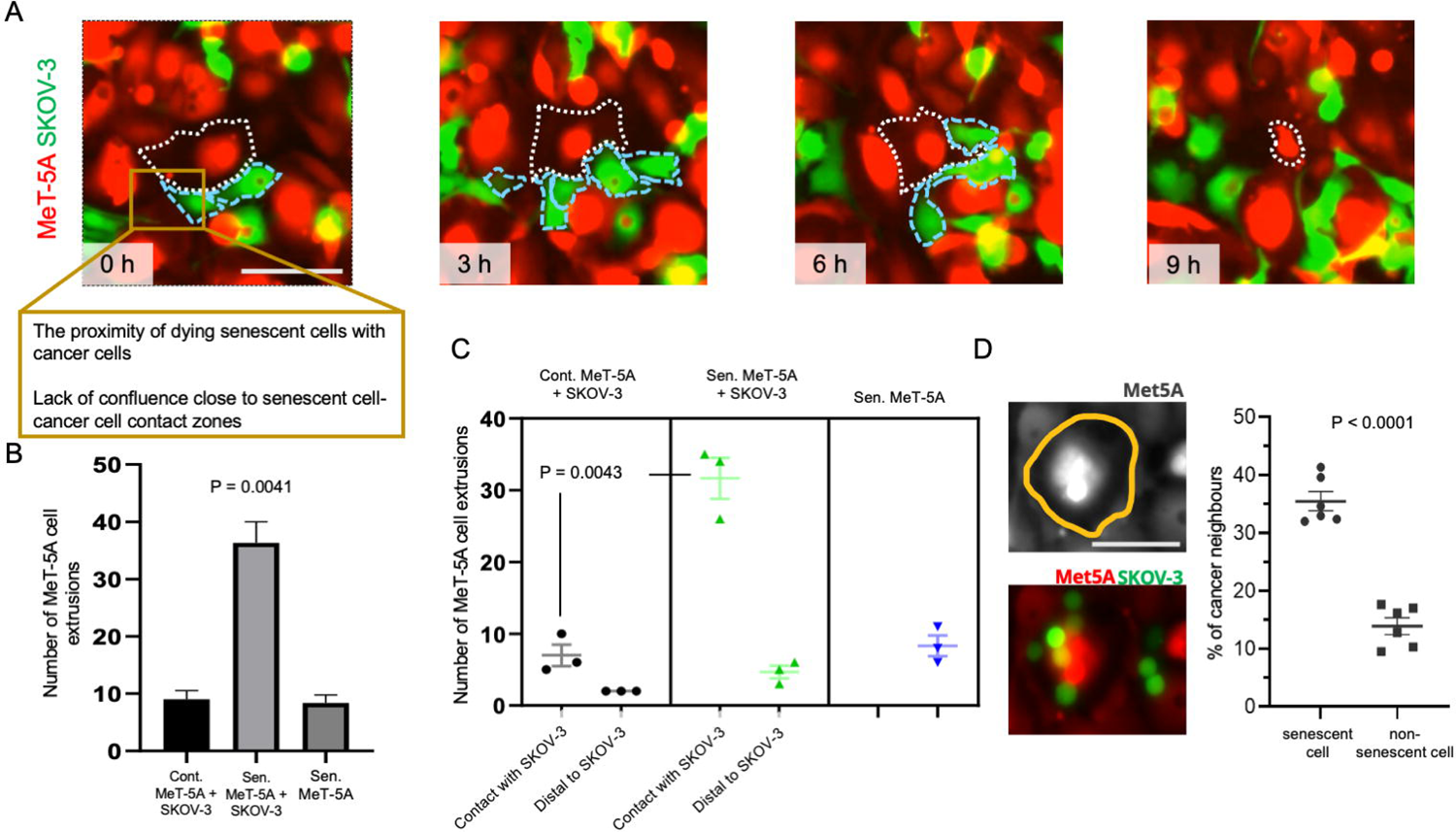
Senescent mesothelia are extruded by several cancer cells localized proximal to them. (A) Representative epifluorescent micrographs of a field imaged at regular time intervals showing the extrusion of senescent RFP-labelled MeT-5A mesothelia by GFP-lavelled SKOV-3 cells as observed in epifluorescence time-lapse videomicroscopy. (B) Bar graph showing the total number of MeT-5A cell extrusions observed in control MeT-5A monolayers with SKOV-3 cells, senescent MeT-5A monolayers with SKOV-3 cells, and senescent MeT-5A monolayers cultured without SKOV-3 cells. (C) Quantitative plot showing quantitatively the number of MeT-5A cell extrusions observed in contact with SKOV-3 cells and distal to SKOV-3 cells in control MeT-5A monolayers and senescent MeT-5A monolayers along with the number of MeT-5A cell extrusions in senescent MeT-5A monolayers without SKOV-3 cells. (D) Representative epifluorescence micrograph showing cancer cells close to a senescent cell (perimeter outlined with yellow) along with the plot showing the percentage of cancerous neighbors for senescent and non-senescent mesothelia within senescent MeT-5A monolayers. The fields are at 10x magnification with a scale bar of 100 μm. The analysis was performed on two biological replicates. The data are presented as mean +/- SEM and significance is obtained using the Brown-Forsythe ANOVA test for (B) and using unpaired parametric t-test with Welch’s correction for (C) and (D).

To further characterize cellular neighborhoods within coculture monolayers, we determined the total number and type of neighbors for senescent and non-senescent mesothelia within cocultures. Due to their larger surface area, the number of neighbors for senescent cells is naturally higher (∼4-5 cells) than for non-senescent cells (∼2-3 cells). Nevertheless, the proportion of neighboring cells being cancer cells was found to be significantly higher for senescent mesothelia than their counterpart non-senescent cells in the same fields (Figure 3D).

Our observation that cancer cells can clear mesothelia independent of cell crowding effects suggested that confluence dependent cell extrusion may not be operative here. This surmise was reinforced by closer localization of cancer cells to senescent mesothelia. To shed light on the intercellular interactions that allow confluence-agnostic senescent cell shedding, we employed a multiscale virtual tissue simulation approach using the Cellular Potts Model (CPM)-based computational framework Compucell3D. A monolayer consisting of senescent, non-senescent, and cancer cells was simulated initially, incorporating differences in their cell proliferation rates and relative cell size. The simulation period in Monte-Carlo steps (MCS) was calibrated with experiments by comparing the growth of cancer cells in the experiments to the growth rate of cancer cells in simulations. The size and number of each cell population was quantified based on their initial starting conditions in experiments. Our first model (Figure 4Bi; video S4) incorporated no other interactive rule. It was, therefore, not surprising that the first model resulted in a confluent monolayer without any colocalization of cancer cells and senescent cells and without any clearance of the latter. Based on our observations in Figure 3A, we incorporated an additional rule (Model 2, Figure 4Bii; video S5) wherein clearance of senescent cells could occur if they share >= 55% of their perimeter with cancer cells. This did result in the extrusion of senescent cells, but only after cancer cells had proliferated enough to create a confluent cocultured monolayer. Therefore, clearance occurred much after confluence had occurred. This led us to hypothesize that an effective attractive force may exist between senescent and cancer cells, driving them to localize close to each other (consistent with our experimental observations of their proximity). Solely implementing such an attractive force (through decreasing the contact energy of adhesion between the senescent and cancer cells; CE_new_ = 15, while CE_old_ = 30) was insufficient for mesothelial clearance (Model 3, Figure 4Biii; video S6). However, the simultaneous implementation of adhesive attraction between senescent and cancer cells and the rule for extrusion (Model 4, Figure 4Biv; video S7) brought about clearance even when the monolayers were sub-confluent, similar to what we observed in experiments (as shown in Figures 4C and 4D). We next asked whether senescent cells secreted any attractive substances that could bring about a closer localization of cancer cells around senescent cells.

**Figure 4:**
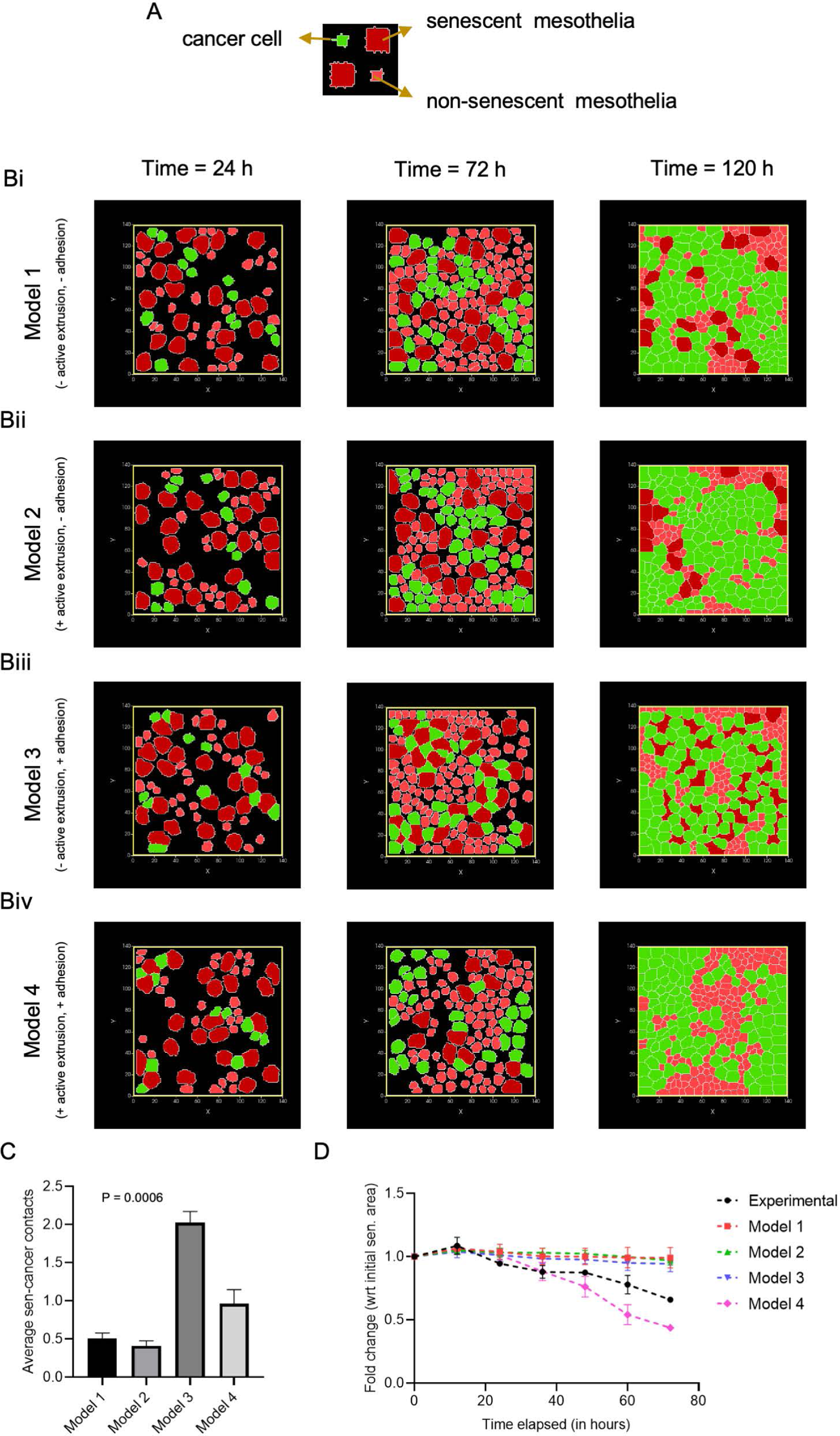
Multicell computational simulations predict the active killing of senescent mesothelial cells by ovarian cancer cells at sub-confluent conditions. (A) The color code of virtual cells used in the simulations. (B) Representative screenshots of cell fields at 24 hours (1000 MCS), 72 hours (3000 MCS), and 120 hours (5000 MCS) for Model 1: no extrusion rule implemented and contact energy (CE) values = 30 (can be seen in video S4) (i), Model 2: extrusion rule and CE = 30 (can be seen in video S5) (ii), Model 3: no extrusion rule and CE = 15 (can be seen in video S6) (iii), and Model 4: extrusion rule and CE = 15 (can be seen in video S7) (iv). (C) bar graph depicting significantly higher senescent cell to cancer cell contacts for Model 4; extrusion rule and CE = 15. (D) Plot comparing senescent mesothelial clearance by ovarian cancer in experiments to the four models used in the study. The data are presented as mean +/- SD, and significance is obtained using the Brown-Forsythe ANOVA test for (C). Model 3 was removed from the statistics in (C) as senescent cells did not undergo clearance at all. The total duration of the simulation was 192 hours (8000 MCS) with a similar initial layout for all four models. The simulations were performed five times for each model.

### A senescent-associated matrisomal phenotype (SAMP) provides a better substratum for the adhesion and spread of ovarian cancer cells

Senescent cells are well known to secrete a gamut of soluble cytokines that may regulate the attraction and proliferation of cells surrounding them, a phenomenon referred to as senescence-associated secretory phenotype ^21, 22^. To verify if the secretome of senescent mesothelia could exert such an effect on ovarian cancer cells, we cultivated SKOV-3 cells in the conditioned medium from senescent (and control non-senescent) cells for 36 hours (Figure S7Ai). We did not observe any significant difference between the proliferation of SKOV-3 cells with conditioned media from senescent and non-senescent mesothelia (Figure S7Aii). We then extended our investigation to the extracellular matrix secreted by senescent cells to check the proliferation of the adhered SKOV-3 cells (Figure S7Bi). We did not observe a significant change in the proliferation of SKOV-3 cells in the senescent and non-senescent extracellular matrices (Figure S7Bii).

However, to our surprise, SKOV-3 cells occupied a larger proportion of decellularized senescent cell ECM than non-senescent controls (Figure 5A). Upon quantification of cell number and size, we found that the greater occupancy was due to the increased number and surface area of adhered SKOV-3 on the decellularized senescent mesothelial ECM, compared with controls, respectively. (Figure 5Bi - ii). Moreover, migrating SKOV-3 cells showed higher velocity and more persistent motion on senescent mesothelial ECM than non-senescent controls (Figure 5C i-ii; can be seen in videos S8 and S9). In support of our observations on SKOV-3, OVCAR-3 cells did not show any difference in proliferation with conditioned media from senescent and non-senescent mesothelia (Figure S8A). We observed no difference in proliferation when grown on decellularized senescent versus non-senescent ECM (Figure S8B); however, we found that a significantly higher number of OVCAR-3 cells adhered to and spread to a greater extent on senescent mesothelial ECM (Figure S9A and S9B i – ii). We then asked what proteins within senescent ECM may be responsible for their ability to enhance adhesion and motility of cancer cells.

**Figure 5:**
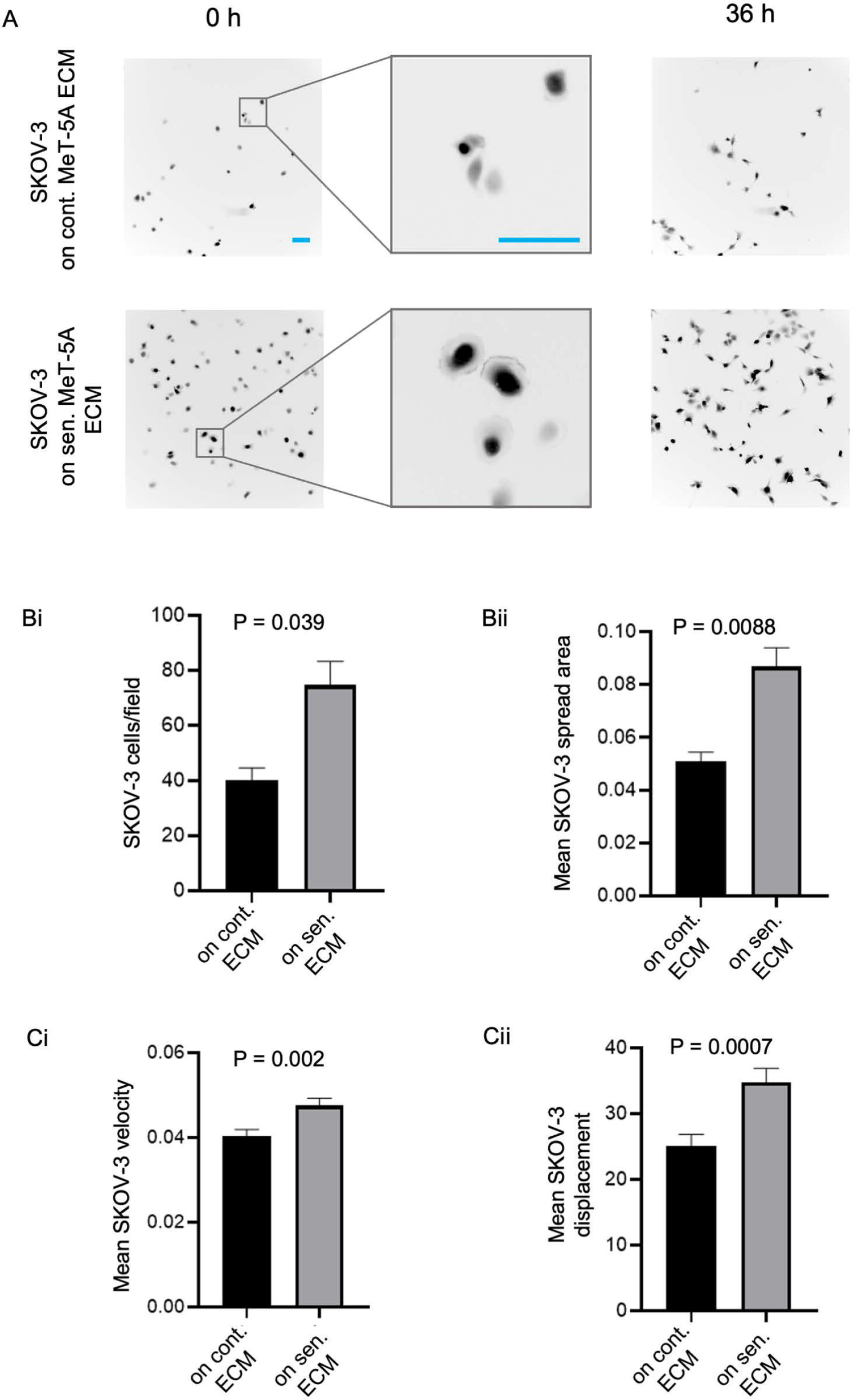
Effect of extracellular matrix secreted by control and senescent MeT-5A monolayers on SKOV-3 adhesion and migration. (A) Representative epifluorescent micrographs of adhered SKOV-3 cells on extracellular matrix secreted by control and senescent MeT-5A monolayers along with insets showing the spread of SKOV-3 cells on senescent extracellular matrix. (B) (i) Bar graph showing the difference in the number of adhered SKOV-3 cells per field on the extracellular matrix secreted by control and senescent MeT-5A monolayers. (ii) Bar graph showing mean SKOV-3 spread area on the extracellular matrix secreted by control and senescent MeT-5A monolayers. (C) (i) Bar graph showing mean velocity of SKOV-3 migration on the extracellular matrix secreted by control and senescent MeT-5A monolayers, 24 hours post adhesion. (ii) Bar graph showing mean SKOV-3 displacement on the extracellular matrix secreted by control and senescent MeT-5A monolayers, 24-hours post adhesion (can be seen in videos S8 and S9). The fields are at 10x magnification with a scale bar of 100 μm. The experiments were performed in triplicates. The data are presented as mean +/- SEM and significance is obtained using an unpaired parametric t-test with Welch’s correction.

### Senescent mesothelial cells secrete higher levels of fibronectin, laminins, and hyaluronic acid

Cells adhere to the extracellular matrix through interactions mediated by their cell surface receptors. Adhesion is predominantly mediated by interactions between integrins on the cell surface and laminin ^23^ and fibronectin ^24, 25^ on the substrata, but also through interactions between cell surface CD44 and hyaluronan ^26, 27^. Therefore, we first performed immunofluorescence on senescent and control MeT-5A monolayers to characterize the differences in levels of laminins and fibronectin within the two monolayers. We observed significantly higher levels of fibronectin (Figure 6Ai) and pan laminin (Figure 6Bi) in senescent MeT-5A monolayers compared to control counterparts. Quantitative RT-PCR confirmed the same and showed a 1.8-fold change in fibronectin expression (Figure 6Aii) and a 4.5-fold change in the expression of laminin B3 (Figure 6Bii) for senescent mesothelia compared with controls. The strong presence of fibronectin in the senescent mesothelial microenvironment was even observed after the culture was decellularized (Figure S10A i – ii), suggesting senescent mesothelia do secrete higher levels of fibronectin and other ECM proteins that constitute the adhesive substratum around themselves. We performed Alcian blue staining on senescent and control MeT-5A monolayers to characterize the differences in the expression of acidic mucins within the two monolayers (Figure 6Ci). We observed significantly higher levels of hyaluronic acid glycosaminoglycans in senescent MeT-5A monolayers than in control counterparts (Figure 6Cii).

**Figure 6:**
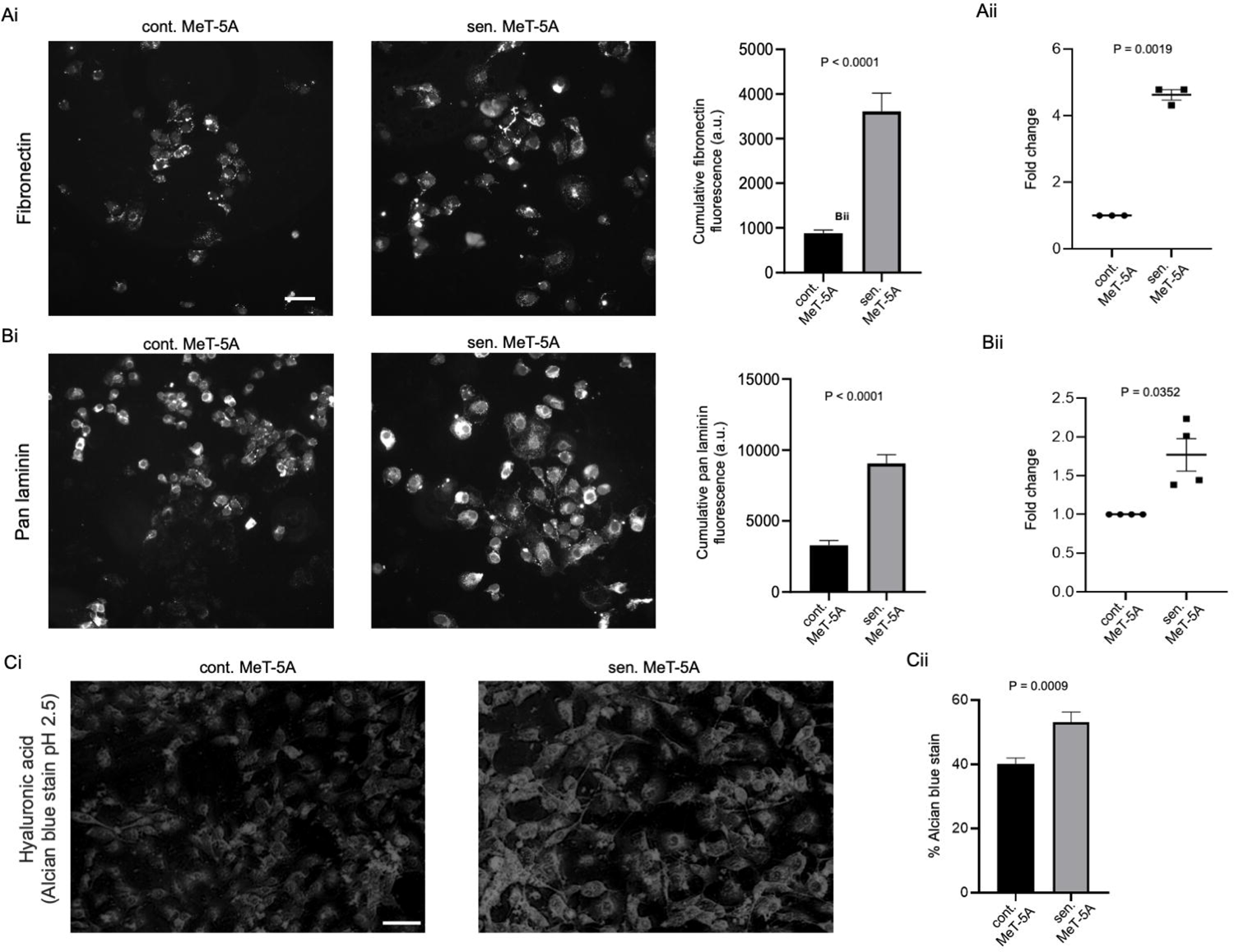
Senescence-associated matrisomal phenotype (SAMP) is characterized by a higher expression of specific extracellular constituents. (A) (i) Representative immunofluorescence micrographs performed for fibronectin on extracellular matrix secreted by control and senescent MeT-5A monolayers. The bar graph shows the cumulative fluorescence intensity of fibronectin within senescent cells compared to non-senescent cells. (ii) qPCR data shows the fold change in fibronectin expression in senescent MeT-5A monolayers compared to control MeT-5A monolayers. (B) (i) Representative immunofluorescence micrographs performed for pan laminin on extracellular matrix secreted by control and senescent MeT-5A monolayers. The bar graph shows the cumulative fluorescence intensity of pan laminin within senescent cells compared to non-senescent cells. (ii) qPCR data shows the fold change in the expression of laminin B3 in senescent MeT-5A monolayers compared to control MeT-5A monolayers. (C) (i) Representative micrographs of alcian blue staining for hyaluronic acid on control and senescent MeT-5A monolayers. (ii) Bar graph showing alcian blue staining for hyaluronic acid within senescent cells compared to non-senescent cells. The fields are at 20x magnification with a scale bar of 50 μm for (A) and (B) and at 10x magnification with a scale bar of 100 μm for (C). The experiments were performed in triplicates. The data are presented as mean +/- SEM and significance is obtained using an unpaired parametric t-test with Welch’s correction.

### Increased expression of fibronectin and laminins aid in the greater adhesion of ovarian cancer cells on senescent extracellular matrix

Ovarian cancer cells are known to bind hyaluronic acid via the CD44 receptor that aids in the adhesion of cancer cells to the peritonea ^28^. Ovarian cancer cells are known to adhere to the peritoneal mesothelia by canonical interactions between beta-1 integrins and fibronectin ^29^. Therefore, to study which of these may be contributing canonically to cell adhesion, we used a soluble cyclic RGD peptide in adhesion assays on senescent and non-senescent matrices as an allosteric inhibitor to hinder the canonical fibronectin- and laminin-integrin binding (Figure 7A i – ii). As expected, with the addition of cyclic RGD peptide, we observed a decline in the number of adhered SKOV-3 cells on the senescent extracellular matrix (Figure 7B). In fact, at 20 μg/mL of cyclic RGD peptide, the adhesion of SKOV-3 cells on senescent extracellular matrix was even lower than that on the non-senescent extracellular matrix (Figure 7C).

**Figure 7:**
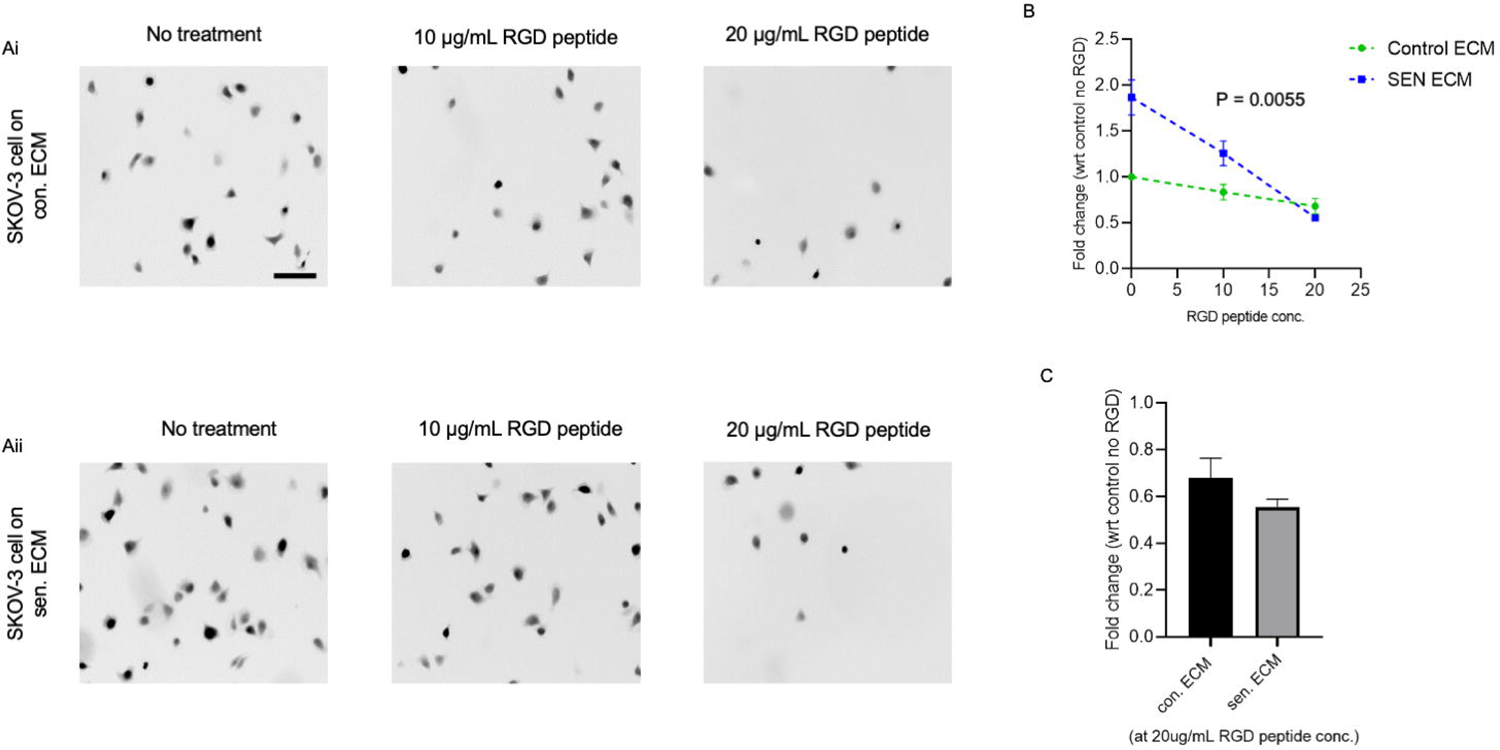
Greater adhesion of ovarian cancer cells on the senescent extracellular matrix is decreased using a synthetic RGD peptide. (A) (i) Representative micrographs of adhered SKOV-3 cells on control mesothelial ECM treated without or with 10- and 20-μg/mL of cyclic RGD peptide. (ii) Representative micrographs of adhered SKOV-3 cells on senescent mesothelial ECM treated without and with 10- and 20-μg/mL concentration of cyclic RGD peptide. (B) Graph showing decrease in the number of adhered SKOV-3 cells at increasing concentrations of cyclic RGD peptide for control and senescent mesothelial ECM. (C) Bar graph comparing the fold change of the number of adhered SKOV-3 cells on control and senescent mesothelial ECM at 20 μg/mL concentration of cyclic RGD peptide. The fields are shown at 10x magnification with a scale bar of 100 μm. The experiments were repeated four times. The data are presented as mean +/- SEM and significance is obtained using an unpaired parametric t-test with Welch’s correction.

## Discussion

Aging and senescence have been proposed to contribute to a higher incidence of epithelial ovarian cancer ^30, 31^. In fact, secretory cytokines of aged mesothelia have, through elegant experiments, been shown to account for a two-fold increase in peritoneal metastasis in aged patients ^17^. However, cellular senescence is not just a consequence of multiple rounds of replication associated with aging, but also to exposure of DNA-adducing chemicals such as doxorubicin and carboplatin. Therefore, it is imperative to investigate how such therapy-induced senescence (TIS) may exacerbate the progression of ovarian cancer, since it may impact decisions by clinicians on the administrative regimen of such drugs ^32, 33^.

Our investigations led us to observe that senescent mesothelia enhanced not just the adhesion of suspended ovarian cancer cells but also their proliferation and movement once the cells attached and were part of the monolayer. The proliferation is likely caused by the enhanced clearance of senescent mesothelia as was evidenced through time lapse imaging. Surprisingly, though sencescent mesothelial extrusion was not a consequence of crowding within the population of mesothelial and cancer cells. Imaging revealed that the cancer cells preferentially localized proximal to senescent mesothelia and cleared them even under subconfluent conditions. Computer simulations have been shown to be useful in predicting the diversities in phenotype of interacting cells with distinct traits ^34–37^. We used a Cellular Potts Model-based simulation framework to explore the interaction between senescent, non-senescent and transformed cells. Our computational results indicate that to achieve the rate of clearance of senescent cells observed in experiments required both a mechanism of ‘active extrusion’ of senescent cells by cancer cells *and* high adhesive interactions between senescent mesothelia and cancer cells. None of these mechanisms were sufficient without the other. To our surprise, we found that the senescent mesothelia secreted an ECM that was preferentially adhesive to cancer cells (in comparison to the non-senescent mesothelial ECM). Senescent mesothelial ECM was shown to be composed of higher levels of fibronectin, laminin and hyaluronic acid, all of which are well known adhesive ECM constituents. Cancer cells not just adhered faster on senescent ECM, they showed increased spreading. In addition, both their velocity and persistence of migration direction were greater on senescent ECM, suggesting a coupling between these two traits (as has been proposed for mesenchymal migration) ^38, 39^. We define these cellular behaviors collectively as senescence-associated matrisomal phenotype (or SAMP, in semantic consonance with the well-known senescence associated secretory phenotype (SASP) ^40, 41^).

Ovarian cancer cells that were treated with conditioned media from senescent mesothelia did not a significant difference in adhesion or proliferation leading us to surmise that the effects of mesothelial senescence on cancer cell phenotype are predominantly matrisomal and not secretory. However, the time course of these experiments is 24 hours, disallowing a rigorous discounting of secretory effects especially on cell proliferation. Experiments on non-senescent mesothelia show their clearance by cancer cells is dependent on forces generated by myosin ^42^. Although not explored in this manuscript, the proximal localization of mesenchymal cancer cells near senescent mesothelia before the latter are extruded suggests a similar mechanism is at play here, and will be experimentally probed in the future.

Prominent experiments describing universal coupling of speed and persistence (UCSP) as being canonical to motile cells have been performed on fibronectin-coated substrata ^39, 43^. This led us to probe and subsequently demonstrate that ovarian cancer cells adhere strongly to senescent ECM likely through adhesive interactions between integrins and RGD-containing ECM glycoproteins such as laminins- and fibronectin. Which tertiary laminin (or any other novel ECM protein) is specifically involved remains to be discovered and will likely be revealed through quantitative mass spectrometric analyses on the senescent and non-senescent matrisomes, followed by validation through perturbations in expression of identified target proteins. In our study, not just ECM proteins, but the non-conjugated glycosaminoglycan hyaluronan is also upregulated in senescent mesothelia. In this manuscript, we have not tested its role in cancer cell adhesion. Using genetic perturbation and pharmacological inhibition, we aim to probe how interactions likely with CD44 may also potentiate adhesion of cancer cells on senescent peritoneal environments.

In spite of the stated limitations of our experimental study, it is evident that the accumulation of senescent mesothelial cells as an unintended consequence of chemotherapeutic exposure aids ovarian cancer progression. The supplementation of chemotherapy with senolytics or targeted inhibitors of cell-ECM adhesion may help mitigate such prometastatic effects ^44, 45^. On a broader note, these observations argue for a broader and holistic approach to cancer management, one which targets not just cancer cells but their untransformed cellular allies within the tumor microenvironment.

## Materials and Methods

### Cell culture

MeT-5A cells purchased from ATCC were maintained in Medium 199 media (HiMedia; AL014A) along with 10 % fetal bovine serum (Gibco; 10270), SKOV-3 cells were maintained in McCoy’s 5A media (HiMedia; AL057A) along with 10 % fetal bovine serum (Gibco; 10270), and OVCAR-3 cells were maintained in RPMI-1640 media (HiMedia; AL162A) along with 20% fetal bovine serum (Gibco; 10270) in a 5% carbon dioxide, 37° C temperature humidified incubator.

### Senescence induction

Non-senescent MeT-5A cells were seeded in 60 mm dishes from an early passage of healthy MeT-5A cells growing and allowed to adhere for 18 hours. Then they were treated with 100 nM of doxorubicin (Cayman chemicals; 15007) for 48 hours ^46, 47^. The control sample was added with a proportionate volume of DMSO to negate the effect of solvent on MeT-5A cells present in the treated samples. The control and treated samples were cultured in drug-free media for 24 hours and then checked for senescence ^47^.

BALB/c female mice (4-6 weeks old) were sacrificed for ex-vivo adhesion experiments. Mice were euthanized by cervical dislocation and confirmed dead by cessation of respiration. The peritoneal membrane was surgically dissected and cut into two parts and cultured in Medium 199 media (HiMedia; AL014A) along with 10% fetal bovine serum (Gibco; 10270) with and without 100nM of doxorubicin to generate senescent and non-senescent murine peritonea, respectively. Gfp-labeled SKOV-3 were suspended on top of the murine peritonea and were allowed to adhere for 30 minutes. The murine peritonea were washed with 1 x PBS and imaged by using an Olympus IX83 inverted epifluorescence microscope at 10X and 4X magnification. The images were analyzed and the number of cells adhered to the matrices was counted using the “analyze particles” feature in ImageJ. Murine cells were isolated from senescent and non-senescent peritonea by incubating them in 0.25% trypsin solution for 8 mins, followed by streaking over the tissue with a tip to dislodge the cells from the tissue. Extracted cells allowed to adhere for 48 hours and used for morphological examination for senescence.

### Cell staining

MeT-5A cells were directly seeded in an eight-well chambered cover glass for staining the DNA and F-actin. Monolayers of senescent and non-senescent MeT-5A cells were fixed using 4 % formaldehyde (24005; Thermo Fisher Scientific) at 4°C for 20 min. After fixation, the cells were taken for further processing or stored in 1× PBS at 4°C. Permeabilization was achieved using 0.5% Triton X-100 (MB031; HiMedia) for 90 mins at RT. Alexa Fluor 488 Phalloidin (Invitrogen; A12379) dissolved in 0.1% Triton X-100 was incubated overnight at 4°C in dark. This was followed by washes using 0.1% Triton X-100 in PBS (5 mins × 3). Samples were incubated with DAPI (D1306; Thermo Fisher Scientific) dissolved in 0.1% Triton X-100 for 10 min in dark at RT. Subsequent processing was carried out in the dark. These included washes using 0.1% Triton X-100 in PBS (5 mins × 2). The images were captured at 10x magnification using an Olympus IX83 inverted epifluorescence microscope and an Olympus IX83 inverted fluorescence microscope fitted with Aurox structured illumination spinning disk setup. Images were processed and analyzed using ImageJ.

### Cytochemical detection of SA-**β**-Gal

MeT-5A cells were fixed with 0.2 % glutaraldehyde in PBS for 15 min at room temperature (RT). The cells were washed twice with PBS and incubated overnight in fresh SA-β-gal staining solution containing 1 mg/ml X-Gal, 5 mM potassium ferrocyanide, 5 mM potassium ferricyanide, 150 mM NaCl, 2 mM MgCl2, and 0.1 m phosphate buffer, pH 6.0 in darkness at 37° C without CO_2_ ^48^. Upon development of the bluish-green stain post-incubation period, images were visualized using a phase-contrast microscope and captured on a Magcam DC camera using MagVision software. Quantitative analysis was performed on the images using ImageJ.

### Mesothelial clearance assay

Non-senescent and senescent RFP-labelled MeT-5A cells were trypsinized and seeded (40,000 in number) in eight well-chambered cover glasses. Sub-confluent monolayers were allowed to form for the next 24 hours and GFP-labelled SKOV-3 cells were added to the non-senescent and senescent monolayers. The assay was performed with Medium 199 media (HiMedia, AL014A) along with 10% fetal bovine serum (Gibco, 10270). Time-lapse imaging was subsequently performed for 72 h right after the addition of SKOV-3 cells at 30 min intervals using a Tokai Hit stage-top incubator with image acquisition through an Orca Flash LT plus camera (Hamamatsu) on an Olympus IX73 microscope. The videos were stitched using ImageJ and the same was used to calculate the change in the area of MeT-5A monolayers and SKOV-3 cells in the same fields (shown in figure S11).

### Quantitative Real-Time PCR

Quantitative real-time PCR was performed for common matrisomal genes, where GAPDH was used as an internal control for the normalization of RT qPCR data. Total RNA was isolated using RNAiso Plus from MeT-5A monolayers, after which 1 μg of total RNA was reverse transcribed to cDNA using Verso cDNA Synthesis kit as per the manufacturer’s protocol (AB-1453; Thermo Fisher Scientific). Real-time qPCR was performed on QIAGEN Rotor-Gene Q System (QIAGEN) using a standard two-step amplification protocol followed by a melting curve analysis. The amplification reaction mixture (total volume of 10 μL) contained 10 ng of cDNA, 5 μL 2× DyNAmo Flash SYBER Green master mix, and 0.25 μM of the appropriate forward and reverse primer. Primer sequences of GAPDH and matrisomal genes are mentioned in the Table S1. Relative gene expression was calculated using the comparative C_t_ method, and gene expression was normalized to non-senescent MeT-5A cells. Appropriate no template and no RT controls were included in each experiment. All the samples were analyzed in triplicates and repeated at least three times independently.

### Immunostaining and image acquisition

MeT-5A cells were directly seeded in an eight-well chambered cover glass for immunostaining. Non-senescent and senescent MeT-5A cells were fixed using 4 % formaldehyde (24005; Thermo Fisher Scientific) at 4°C for 20 min. After fixation, the cells were taken for further processing or stored in 1× PBS at 4°C. Permeabilization was achieved using 0.5% Triton X-100 (MB031; HiMedia) for 90 mins at RT. Blocking was achieved using PBS with 0.1% Triton X-100 and BSA (MB083; HiMedia) for 45 min at RT. Primary antibody incubation was carried out overnight at 4°C. This was followed by washes using 0.1% TritonX-100 in PBS (5 min × 3). Secondary antibodies and Alexa Fluor 488/568 phalloidin (Invitrogen; A12379; A12380) were incubated with the sample at RT for 2 h under dark conditions. DAPI (D1306; Thermo Fisher Scientific) was added to the samples and washed away after 15 min. Subsequent processing was carried out in the dark. These included washes using 0.1% Triton X-100 in PBS (5 mins × 3). Images were captured at 10x magnification using an Olympus IX83 inverted epifluorescence microscope. Images were processed and analyzed using ImageJ. The antibodies used in our studies are against pan-Laminin (ab11575) and fibronectin (E5H6X). Negative controls in each case were through the omission of the primary antibody.

### Alcian blue staining

MeT-5A cells were directly seeded in an eight-well chambered cover glass for alcian blue staining. Non-senescent and senescent MeT-5A cells were fixed using 4 % formaldehyde (24005; Thermo Fisher Scientific) at 4°C for 20 min. After fixation, the cells were washed three times with 1 x PBS and then taken for further processing or stored at 4°C. While staining for Hyaluronic acid and acid mucins, the MeT-5A monolayers were incubated with 3% acetic acid solution for three minutes and then with the alcian blue solution (1% w/V alcian blue, 8GX (HiMedia, TC359-10G) in 3% acetic acid solution at 2.5 pH) for 30 minutes at 37°C. Perform three 1 x PBS washes for 5 minutes each. Upon development of the bluish-green stain, images were visualized using a phase-contrast microscope and captured on a Magcam DC camera using MagVision software. Quantitative analysis was performed on the images using ImageJ.

### Conditioned media treatment

The seeded non-senescent MeT-5A cells were made senescent by incubation with 100 nM of doxorubicin (Cayman chemicals; 15007) for 48 hours. The media along with the remaining doxorubicin was removed and the senescent monolayer was washed with 1 x PBS. Senescent monolayers were incubated with doxorubicin-depleted media for another 24 hours and the conditioned media (CM) was collected. Non-senescent MeT-5A monolayers were also incubated with media for 24 hours and the CM was collected. A 1:1 mixture of CM from senescent/non-senescent MeT-5A and Medium 199 media (HiMedia; AL014A) along with 10 % fetal bovine serum (Gibco; 10270) were used for conditioned media growth assay with SKOV-3 cancer cells.

### Senescent and non-senescent matrix preparation

Senescent and non-senescent MeT-5A cells were seeded at high confluency on chamber wells with coverslip bottoms. The cells were allowed to adhere and lay down the extracellular matrix for the next 36 hours. The confluent MeT-5A monolayers were then washed with 1 x PBS twice and decellularized using a 20 mM solution of NH_4_OH in Milli-Q water for 7-10 minutes ^49^. The cellular debris was gently removed by giving 2-3 washes with Milli-Q water. The extracted extracellular matrices laid on the chamber wells were used for adhesion assays.

### Adhesion assay with RGD peptide

The SKOV-3 cells (5000 in number) suspended in Mcoy’s 5A media (HiMedia; AL057A) along with 10 % fetal bovine serum (Gibco; 10270) were added to the decellularized senescent and non-senescent matrices and allowed to adhere on the matrix for 25 mins. For adhesion assay with RGD peptide, the media also had cyclic RGD peptide (Sigma; G1269-1MG) at 10 and 20 μg/mL concentrations. After incubation, the wells were washed with 1 x PBS twice and immediately imaged by using an Olympus IX83 inverted epifluorescence microscope at 10x magnification.

The images were analyzed and the number of cells adhered to the matrices was counted using the “analyze particles” feature in ImageJ. The attached cells were also time-lapsed for 24 hours using a Tokai Hit stage-top incubator with image acquisition through an Orca Flash LT plus camera (Hamamatsu) on an Olympus IX73 microscope. The data were analyzed using a manual tracking plugin in FIJI and Ibidi’s chemotaxis and migration tool to see differences in displacement and velocity of SKOV-3 cells on senescent and non-senescent matrices (can be seen in videos S8 and S9).

### Compucell 3D model for the clearance of senescent mesothelia

Compucell3D (CC3D) is based on the Cellular Potts Model (CPM), also known as the Glazier-Graner-Hogeweg (GGH) model. It has been utilized extensively for computational models of the collective behaviour of cellular structures. The CPM is a lattice-based discrete model in which spatiotemporal development is simulated via an energy minimizing procedure ^34, 50^. Each cell consists of a collection of lattice sites (pixels). Each configuration is associated with an effective energy, or Hamiltonian (H), which is calculated based on properties such as volume, surface area, contact energies, or external properties. Time evolution is simulated via Monte Carlo Steps (MCS) that involve random changes of lattice site occupations and the changes that decrease the energy are more likely than those that increase it. The Hamiltonian of the system is formulated as:

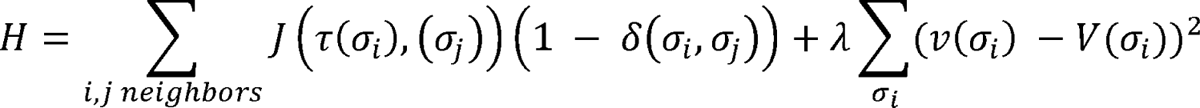

Where *i,j* are lattice sites, σ_*i*_ is the cell at site i, τ(σ) is the cell type of cell σ, J is the coefficient determining the adhesion between two cells of types τ(σ), τ(σ’), δ is the Kronecker delta, v(σ) is the volume of cell σ, V(σ) is the target volume, and λ is a Lagrange multiplier determining the strength of the volume constraint. We utilized a simple two-dimensional model 140*140*1 using a square lattice to find and understand the effect of factors sufficient to bring a considerable decline in the senescent mesothelia area as observed in the mesothelial clearance experiments that were recorded using time-lapse epifluorescence videography. The simulation time was set to 8000 MCS as per the corresponding normalized length of the experimental time-lapse videos. The lattice is composed of the medium, senescent cells (SEN), non-senescent cells (NON_SEN), ovarian cancer cells (CANCER), and a frozen wall that mimics a closed system such as enclosed peritoneal cavity (shown in figure S12). This simple model has four variations with the contact energies between cancer and senescent cells to be either 15 (greater adhesion) or 30 (same as other contact energy combinations) and activation or deactivation of the active extrusion rule (shown in figure S13). The other important parameters used in the model and their input values are available at Table S2. To compare the simulation time to the timescale of senescent mesothelial clearance by ovarian cancer cells, a calibration was performed. The division time of ovarian cancer cells in experiments (SKOV-3 cells) was 18 hours and the division time of ovarian cancer cells in simulations was 750 MCS. This factor was used to calibrate the simulations time to real-time and the comparison between the computational model and senescent mesothelial clearance experiments was made.

### Statistical analysis

The experiments were repeated at least three times and unpaired parametric t-test with Welch’s correction were used for statistical testing unless otherwise stated in the figure legends. The graphs were plotted using GraphPad Prism 8.0.2 and *p-* values < 0.05 was considered statistically significant.

### Code

https://github.com/vivanbharat/MS-Thesis-code

## Acknowledgements

This work was supported by the Wellcome Trust/DBT India Alliance Fellowship/Grant [IA/I/17/2/503312] awarded to RB. It was also supported by the John Templeton Foundation (#62220) (to RB and TG) and by the Department of Biotechnology, India [BT/909 PR26526/GET/119/92/2017] (to RB). DKS acknowledges support from Science and Engineering Research Board (SERB), India (#CRG/2020/000239) and financial support from IISc. BVT acknowledges KVPY for the student scholarship [SX(E-SC)-1711005]. The opinions expressed in this paper are those of the authors and not those of the John Templeton Foundation.

