## Supplementary File for "The senescent mesothelial matrix accentuates colonization by ovarian cancer cells"

S1 A

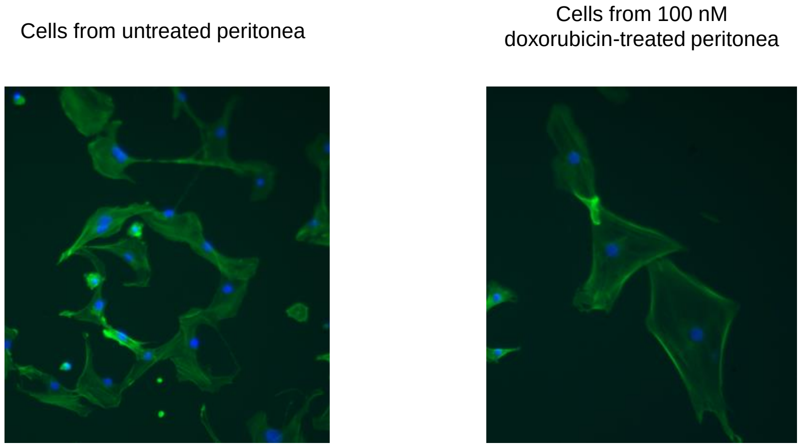


The confirmation of senescent cells acquired from ex-vivo murine peritoneal treated with 100 nM of doxorubicin. (A) The representative snapshots show stained DNA (using DAPI) and F-actin (using phalloidin) of the treated and untreated samples for morphological examination of the non-senescent and senescent cells extract from the murine peritonea. The magnification is 10x, the scale bar is 100 μm, and the experiment was performed in duplicates.

S2 A

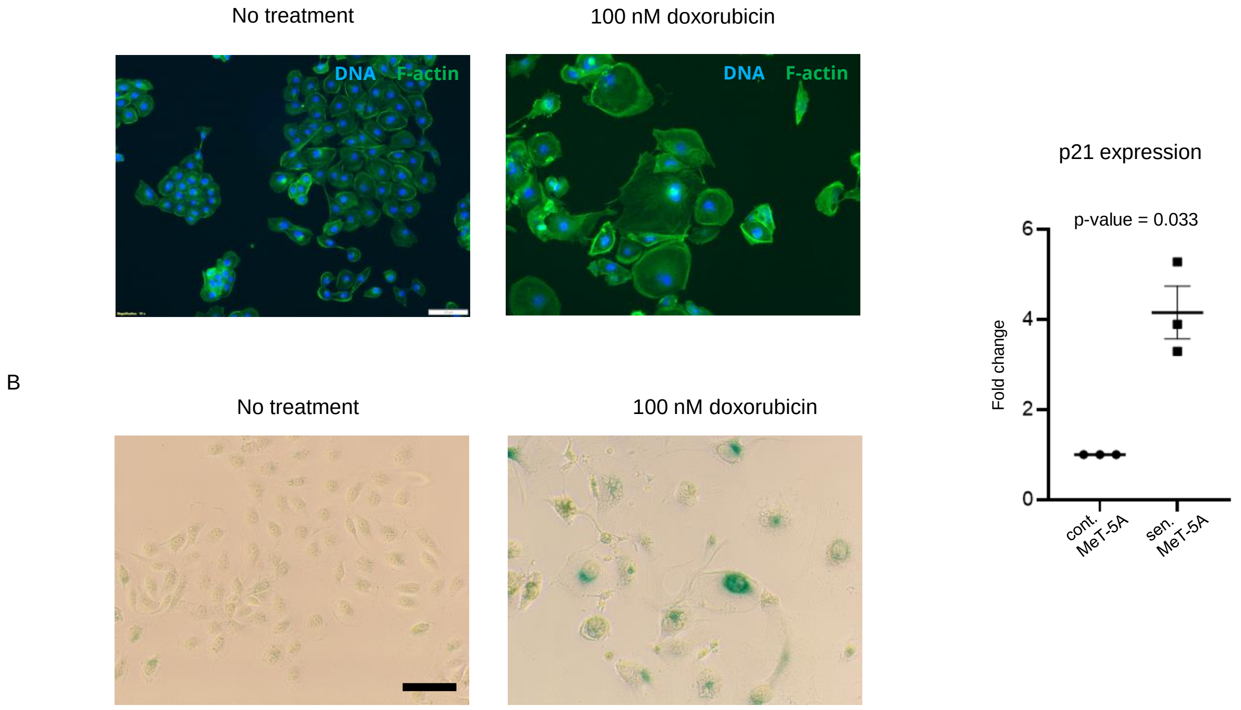


The confirmation of senescent MeT-5A cells upon treatment with doxorubicin.
(A) Staining DNA (using DAPI) and F-actin (using phalloidin) of the treated and untreated samples for morphological examination of the non-senescent and senescent cells, (B) SA-β-gal staining of the treated and untreated samples for the detection of senescent cells with the dichloro-dibromo indigo precipitate (greenish-blue dye) accumulation, (C) qPCR fold change analysis of p21 expression on treated and untreated samples. The magnification is 10x; the scale bar is 100 μm, and p-values were computed using a two-tailed, parametric unpaired t-test with Welch's correction.

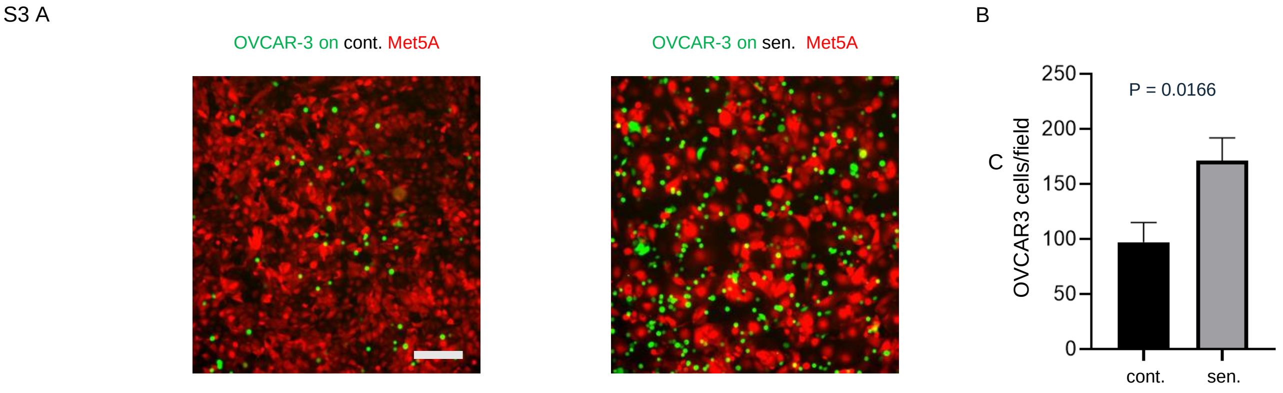


Effect of senescent microenvironment on the adhesivity of ovarian cancer cells.
(A) Representative images show the adhesion of GFP-labelled OVCAR-3 cells on control and senescent MeT-5A monolayers, (B) The bar graph shows the number of OVCAR-3 cells per field for control and senescent MeT-5A monolayers at 10x magnification with a scale bar of 100 µm. The experiments were performed in triplicates. The data are presented as mean +/- SEM and significance is obtained using an unpaired parametric t-test with Welch's correction.

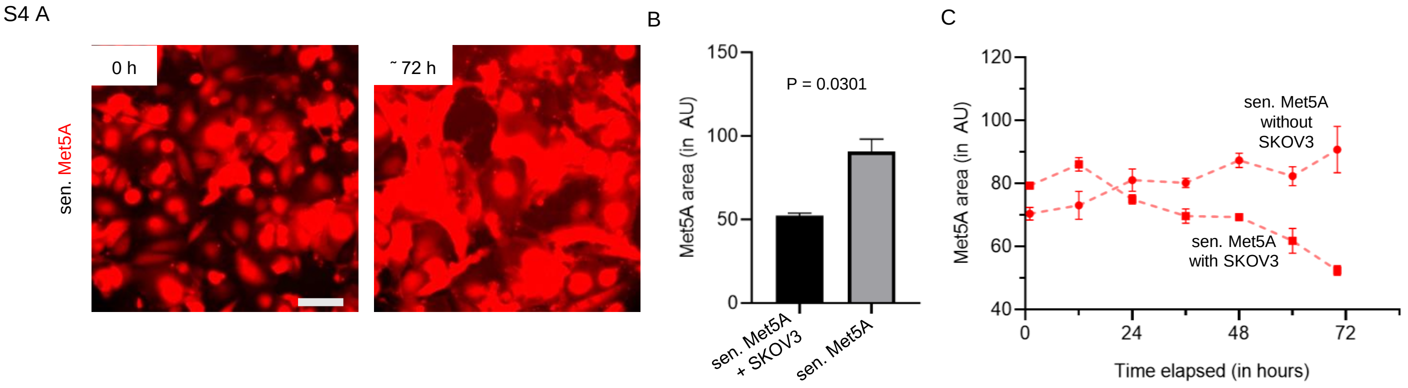


Epifluorescence timelapse videography of mixed cultures of RFP-labelled MeT-5A monolayers with GFP-labelled SKOV-3 cells. (A) Representative images show the area of senescent MeT-5A monolayer without SKOV-3 cells at 0-hour and 72-hour time points (can be seen in video S3). (B) The bar graph compares the area of senescent MeT-5A monolayers with and without SKOV-3 cells at a 72-hour time point. The fields are at 10x magnification with a scale bar of 100 µm. The experiments were performed in triplicates. (C) Quantification of the area of senescent MeT-5A monolayers with and without SKOV-3 cells for 72 hours, plotted with 12-hour intervals. The data are presented as mean +/- SEM and significance is obtained using an unpaired parametric t-test with Welch's correction.

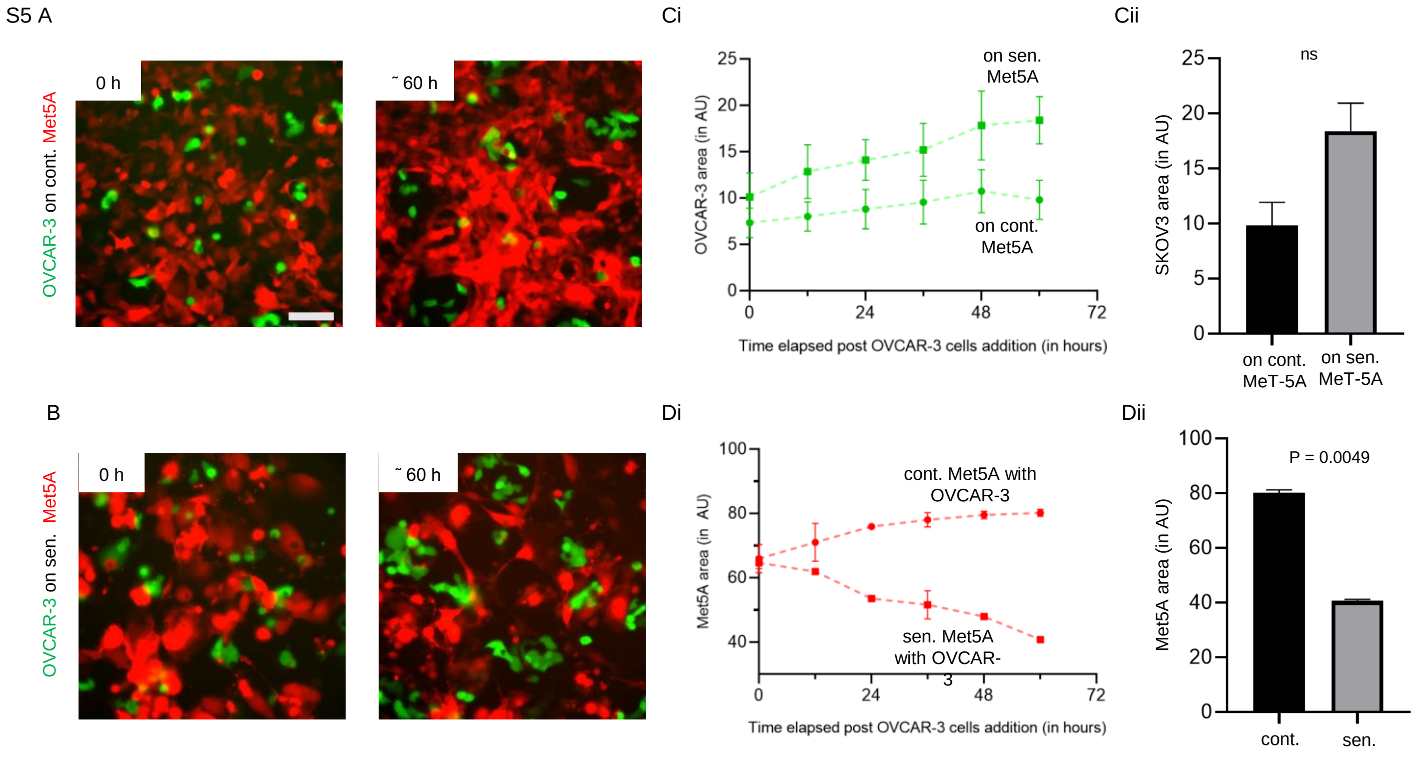


Epifluorescence timelapse videography of mixed cultures of RFP-labelled MeT-5A monolayers with GFP-labelled OVCAR-3 cells. (A) Representative images show the area of control MeT-5A monolayer and OVCAR-3 cells at 0-hour and 60-hour time points. (B) Representative images show the area of senescent MeT-5A monolayer and SKOV-3 cells at 0-hour and 60-hour time points. (C) (i) Quantification of the area occupied by OVCAR-3 cells within control and senescent MeT-5A monolayers for 60 hours, plotted with 12-hour intervals. (ii) The bar graph compares the area of OVCAR-3 cells in the two monolayers at a 60-hour time point. (D) (i) Quantification of the area of control and senescent MeT-5A monolayers for 60 hours, plotted with 12-hour intervals. (ii) The bar graph compares the area of control and senescent MeT-5A monolayers at a 60-hour time point. The fields are at 10x magnification with a scale bar of 100 µm. The experiments were performed in triplicates. The data are presented as mean +/- SEM and significance is obtained using an unpaired parametric t-test with Welch's correction.

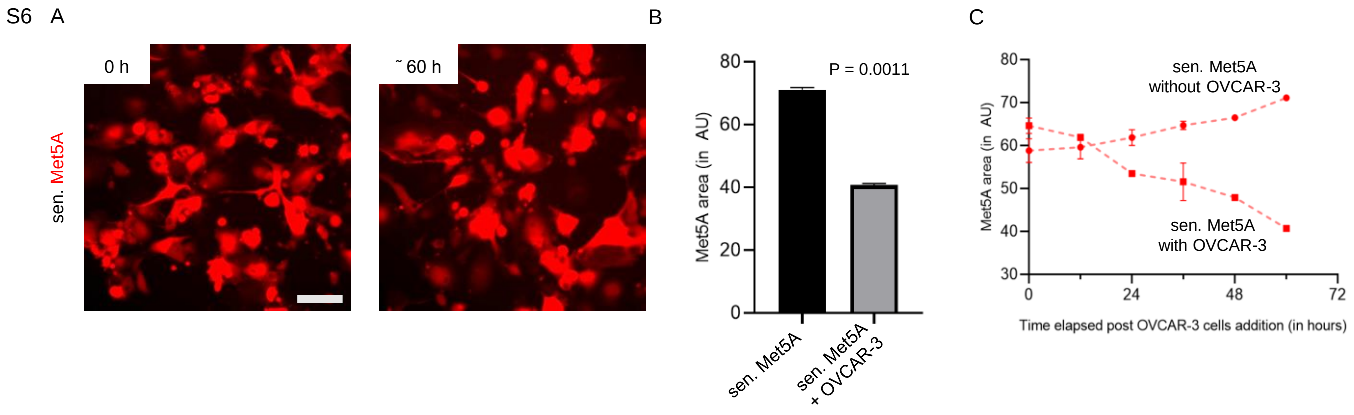


Epifluorescence timelapse videography of mixed cultures of RFP-labelled MeT-5A monolayers with GFP-labelled OVCAR-3 cells. (A) Representative images show the area of senescent MeT-5A monolayer without OVCAR-3 cells at 0-hour and 60-hour time points. (B) The bar graph compares the area of senescent MeT-5A monolayers with and without OVCAR-3 cells at a 72-hour time point. (C) Quantification of the area of senescent MeT-5A monolayers with and without OVCAR-3 cells for 60 hours, plotted with 12-hour intervals. The fields are at 10x magnification with a scale bar of 100 µm. The experiments were performed in triplicates. The data are presented as mean +/- SEM and significance is obtained using an unpaired parametric t-test with Welch's correction.

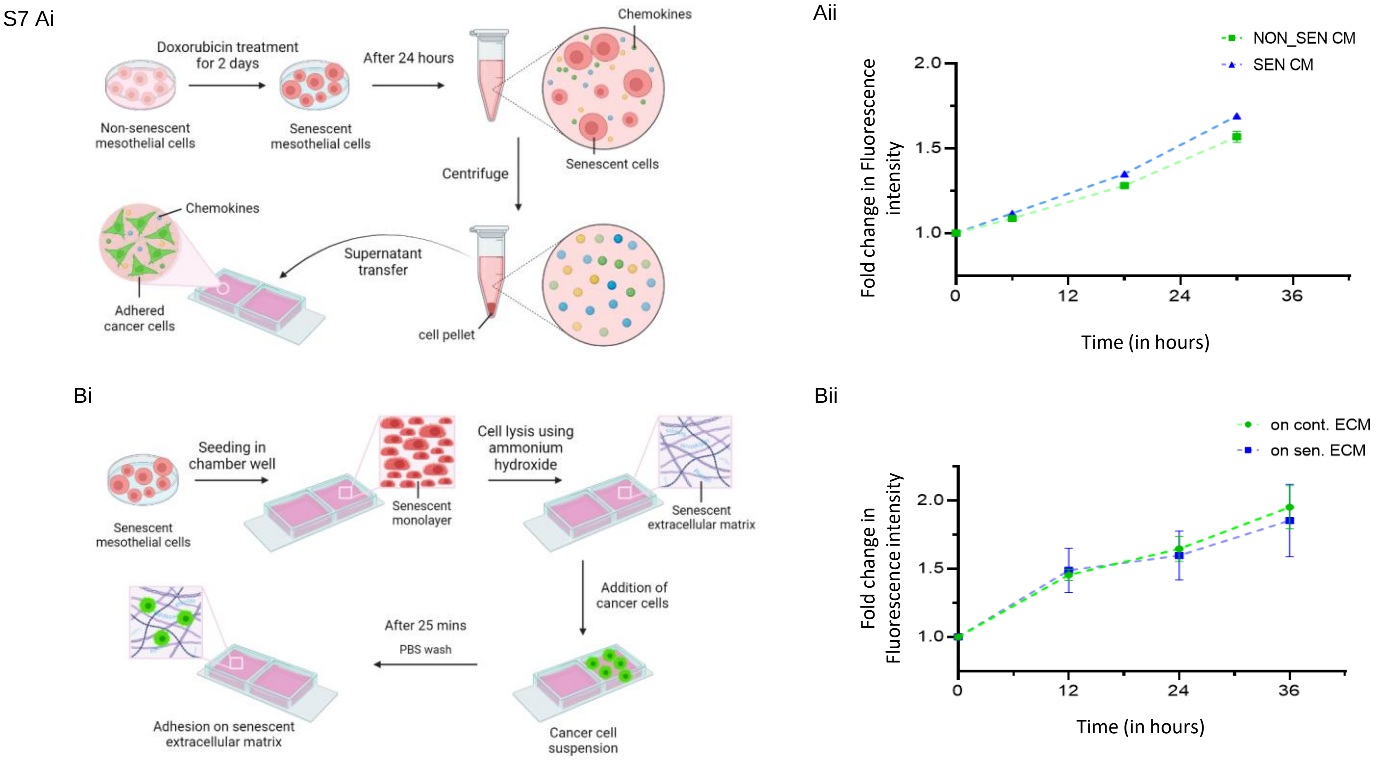


Effect of senescence-associated secretory phenotype (SASP) and extracellular matrix secreted by control and senescent MeT-5A monolayers on the proliferation of SKOV-3 cells. (A) (i) Schematics of proliferation assay performed on SKOV-3 cells using conditioned media from control and senescent MeT-5A monolayers. (ii) Plot shows the fold change in fluorescence intensity of SKOV-3 cells for 30 hours of incubation with conditioned media, plotted with 12-hour intervals. (B) (i) Schematics of adhesion and proliferation assay performed on SKOV-3 cells seeded on extracellular matrix secreted by control and senescent MeT-5A monolayers. (ii) Plot shows the fold change in fluorescence intensity of SKOV-3 cells that adhered to the extracellular matrix for 36 hours, plotted with 12-hour intervals. The experiments were performed in duplicates, and the data are presented as mean +/- SEM.

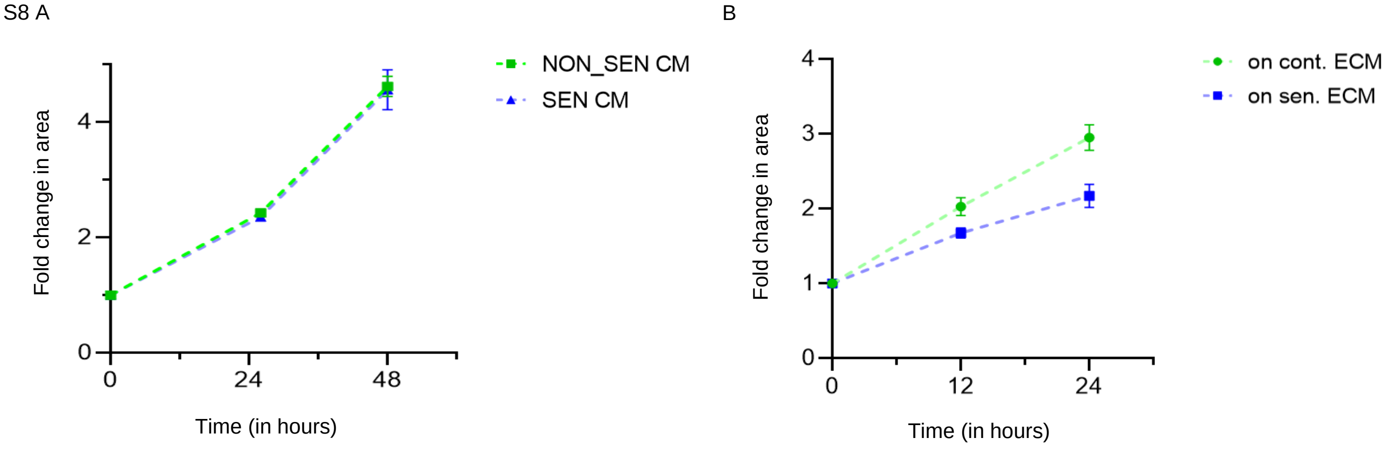


Effect of senescence-associated secretory phenotype (SASP) and extracellular matrix secreted by control and senescent MeT-5A monolayers on the proliferation of OVCAR-3 cells. (A) The plot shows the fold change in fluorescence intensity of OVCAR-3 cells for 48 hours of incubation with conditioned media, plotted with 24-hour intervals. (B) The plot shows the fold change in fluorescence intensity of OVCAR-3 cells that adhered to the extracellular matrix for 24 hours, plotted with 12-hour intervals. The experiments were performed in duplicates, and the data are presented as mean +/- SEM.

S9 A

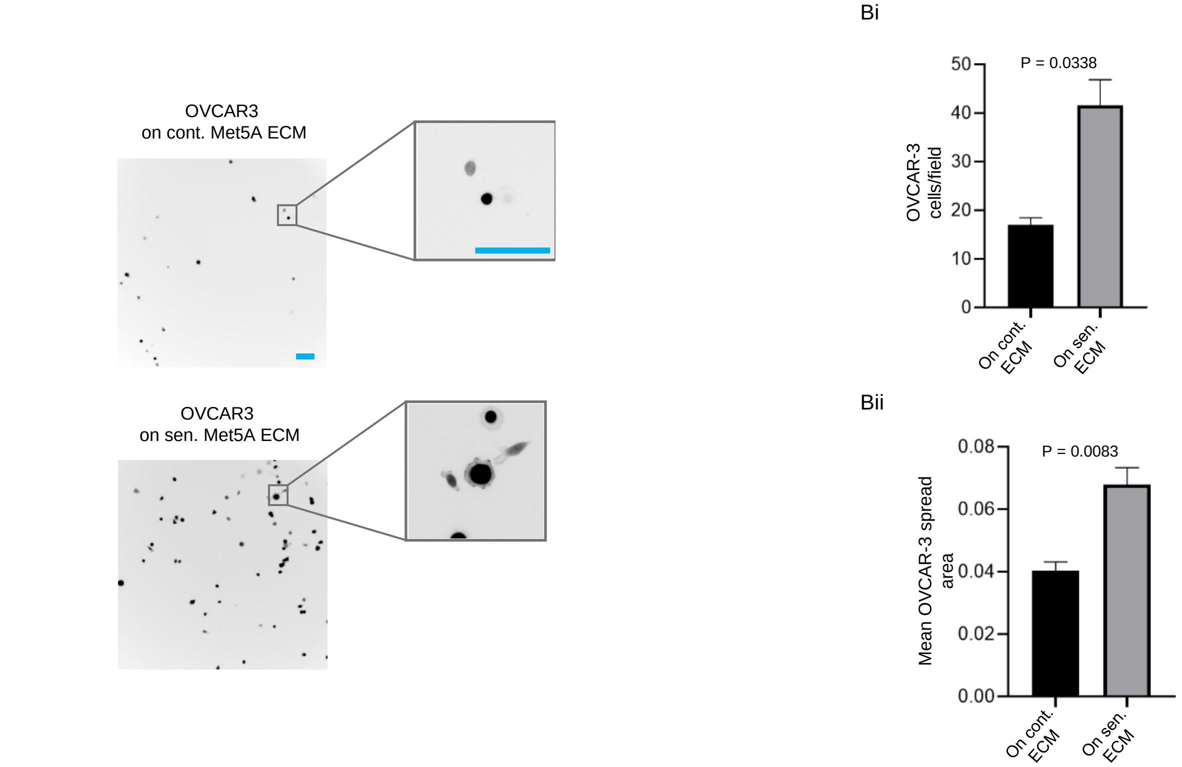


Effect of extracellular matrix secreted by control and senescent MeT-5A monolayers on the adhesivity and spread of OVCAR-3 cells. (A) Representative images of adhered OVCAR-3 cells on extracellular matrix secreted by control and senescent MeT-5A monolayers along with insets showing the spread of OVCAR-3 cells on senescent extracellular matrix. (B) (i) Bar graph shows the number of adhered OVCAR-3 cells per field on the extracellular matrix secreted by control and senescent MeT-5A monolayers. (ii) Bar graph shows the mean OVCAR-3 spread area on the extracellular matrix secreted by control and senescent MeT-5A monolayers. The fields are at 10x magnification with a scale bar of 100 µm. The experiments were performed in triplicates. The data are presented as mean +/- SEM and significance is obtained using an unpaired parametric t-test with Welch's correction.

S10 Ai

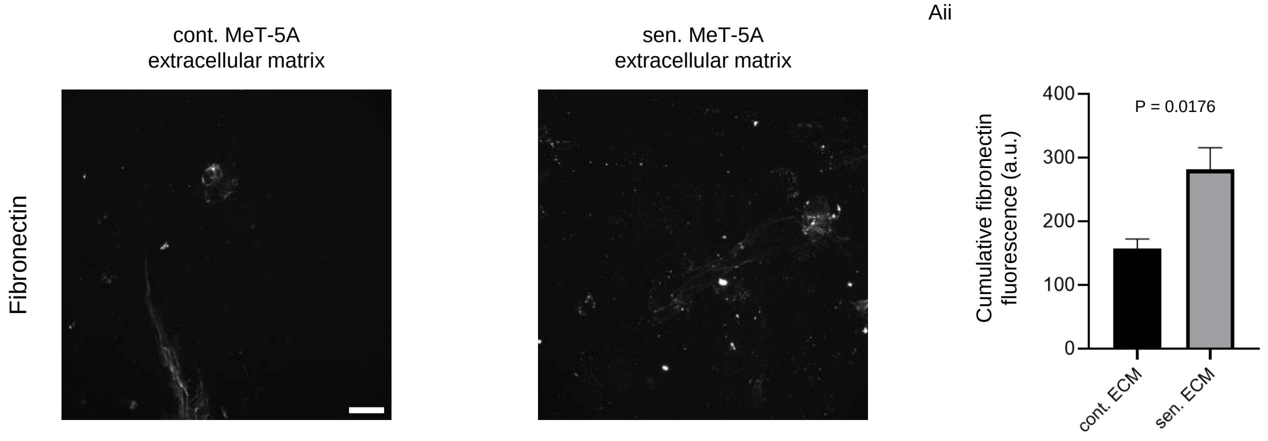


Senescence-associated matrisomal phenotype (SAMP) is characterized by a higher expression of extracellular proteins. (A) (i) Representative images for immunofluorescence performed for fibronectin on decellularized control and senescent MeT-5A monolayers. (ii) The bar graph shows the cumulative fluorescence intensity of fibronectin of entire senescent fields compared to non-senescent fields. The fields are shown at 20x magnification with a scale bar of 50 µm. The experiments were performed in duplicates. The data are presented as mean +/- SEM and significance is obtained using an unpaired parametric t-test with Welch's correction.

S11

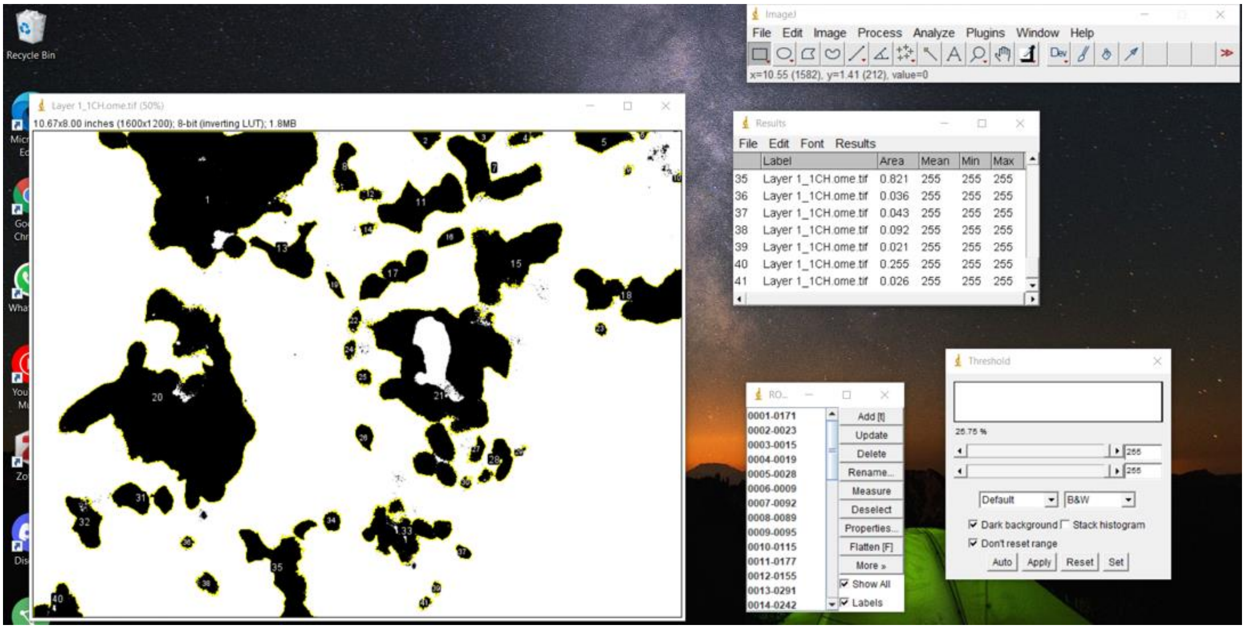


Screenshot showing the analysis performed on images collected from mesothelial clearance assay to calculate the area occupied by MeT-5A and SKOV-3/OVCAR-3 cells in the same field. The highest value of the threshold with a neat background was chosen for generating the mask. The area of cells in the field was measured by summing up the individual particle area and is calculated under the 'Analyze particles' options with size limit = 0.01-Infinity.

S12

| Cell Type Id | Color | Name |
|---|---|---|
| 0 | | Medium |
| 1 | | SEN |
| 2 | | NON_SEN |
| 3 | | CANCER |
| 4 | | WALL |

| Property | Value |
|---|---|
| Revision | 2 |
| Version | 4.3.0 |
| Metadata | Metadata |
| Potts | Potts |
| Plugin | CellType |
| Plugin | Volume |
| Plugin | Surface |
| Plugin | ExternalPotential |
| Plugin | CenterOfMass |
| Plugin | NeighborTracker |
| Plugin | Contact |
| Plugin | Chemotaxis |
| Plugin | Secretion |
| Steppable | DiffusionSolverFE |
| Steppable | UniformInitializer |

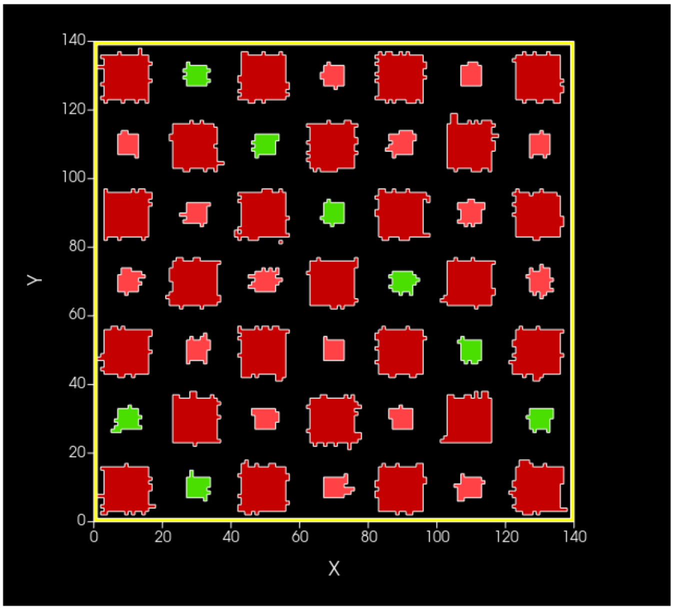


A snapshot of the initial layout of the different cells on the lattice – The picture depicts the initial layout of senescent, non-senescent, and cancer cells in a two-dimensional setup and the incorporated plugins. The relative sizes and the proliferation rates of the three cell types were calibrated from the experiments.

S13

```
common_area_frac = 0.55

for cell in self.cell_list_by_type(self.SEN):
    Avol_SEN += cell.volume/L_SEN
    cancer_common_area = 0
    for neighbor, common_surface_area in self.get_cell_neighbor_data_list(cell):
        if neighbor and neighbor.type == self.CANCER:
            cancer_common_area += common_surface_area
            Average_cancer_neighbours += 1

    # Active clearance rule is impleted in the next three line
    if cancer_common_area/cell.surface > common_area_frac:
        cell.targetVolume = 0
        cell.lambdaVolume = 10000
```

A snapshot of the active clearance code from Twedit++5 – The code shows the implementation of the killing of senescent cells if more than 55% of their perimeter is shared with ovarian cancer cells.

Table S1

| Gene name | Forward primer (5'→3') | Reverse primer (5'→3') |
|---|---|---|
| GAPDH | GGAGCGAGATCCCTCCAAAA | GGCTGTTGTCATACTTCTCATGG |
| p21 | GGAAGACCATGTGGACCTGT | TAGGGCTTCCTCTTGGAGAA |
| Gal-9 | CCTACCTGAGTCCAGCTGTC | GAGGGTTGAAGTGGAAGGCA |
| LamB3 | GCAGCCTCACAACTACTACAG | CCAGGTCTTACCGAAGTCTGA |
| Col6A1 | ACAGTGACGAGGTGGAGATCA | GATAGCGCAGTCGGTGTAGG |
| FN1 | CAAGCCAGATGTCAGAAGC | GGATGGTGCATCAATGGCA |
| LamA3 | CACCGGGATATTTCGGAATC | AGCTGTCGCAATCATCACATT |
| Col18A1 | CAGTGGACACACTTAGCCCTC | GCGGCATTCTCTGGAACTCC |
| FBLN1 | AGAGCTGCGAGTACAGCCT | CGACATCCAAATCTCCGGTCT |
| FBLN-1C | TGACTGGCATCCACAACTGC | GCTTGGAGCACTCCCGATTCT |
| FBLN-1D | TGCCTACCTTCCGCGAGTTC | GCCGTCCATGTAACGCTTGA |
| Gal-1 | TCAAACCTGGAGAGTGCCTT | CACACCTCTGCAACACTTCC |
| Gal-3 | ATGGCAGACAATTTTTCGCTCC | GCCTGTCCAGGATAAGCCC |

Table 1 – RT PCR primer sequence information for the gene expression studies in non-senescent and senescent MeT-5A monolayers

Table S2

| Parameter | Value |
|---|---|
| Temperature ($T_m$) | 20 |
| NeighborOrder | 1 |
| Boundary conditions X (constant derivative) | Min and Max = 0 |
| Boundary conditions Y (constant derivative) | Min and Max = 0 |
| Contact Energies between cell type | |
| Medium-Medium | 10 |
| Medium-SEN | 10 |
| Medium-NON_SEN | 10 |
| Medium-CANCER | 10 |
| SEN-SEN | 30 |
| SEN-NON_SEN | 30 |
| SEN-CANCER | [15, 30] |
| NON_SEN-NON_SEN | 30 |
| NON_SEN-CANCER | 30 |
| CANCER-CANCER | 30 |
| WALL-SEN | 50 |
| WALL-NON_SEN | 50 |
| WALL-CANCER | 50 |
| Volume and other generic parameters for all cell types | |
| Target Volume SEN | 196 |
| $\lambda_{vol}$ SEN | 20 |
| Target Volume NON_SEN | 49 |
| $\lambda_{vol}$ NON_SEN | 20 |
| Target Volume CANCER | 74 |
| $\lambda_{vol}$ CANCER | 20 |
| Probability of cell death on its own | $3.33 \times 10^{-5}$ |
| Common perimeter killing threshold for SEN | 0.55 |
| Maximum MCS | 8100 |

Table 2 – Table showing all the parameters and their possible values used in the computational model.

Video legends:

Video S1 – Coculture of SKOV-3 cells on control MeT-5A monolayer

Video S2 – Coculture of SKOV-3 cells on senescent MeT-5A monolayer

Video S3 – Senescent MeT-5A monolayer without SKOV-3 cells

Video S4 – CC3D senescent mesothelial clearance model 1 (no extrusion rule and CE = 30)

Video S5 – CC3D senescent mesothelial clearance model 2 ( extrusion rule and CE = 30)

Video S6 – CC3D senescent mesothelial clearance model 3 (no extrusion rule and CE = 15)

Video S7 – CC3D senescent mesothelial clearance model 4 (extrusion rule and CE = 15)

Video S8 – The distance and displacement of SKOV-3 cells on control extracellular matrix

Video S9 – The distance and displacement of SKOV-3 cells on senescent extracellular matrix
